# Population-level emulation enables fast Bayesian calibration of cardiac digital twins

**DOI:** 10.64898/2026.11.08.765186

**Authors:** Clara E. Jones, Felix Plappert, Derek J. Bivona, Johane H. Bracamonte, Zarif Hossain, Isabella Aleman, Lyna Luu, Kenneth C. Bilchick, Pim J.A. Oomen

## Abstract

**Purpose:** Patient-specific cardiac digital twins have the potential to guide individualized therapy, but generating these in a routine clinical workflow remains challenging. This is especially true when the twins must carry quantified uncertainty: Bayesian calibration requires thousands of simulations per patient, and the emulators that reduce this cost are typically trained one patient at a time. We aimed to train a single emulator across a patient cohort, so that a new patient can be calibrated in minutes without new simulations.

**Methods:** We trained a population-level Gaussian process emulator whose inputs included a 12-dimensional latent encoding of each patient’s clinical features, so that a single emulator represents the whole cohort. We compared spending simulation budgets on resampling the real cohort with augmenting the chort with digital chimeras, synthetic patients interpolated between real patients. Seven parameters of a closed-loop heart and circulation model were calibrated by Markov chain Monte Carlo to ten clinical biomarkers in 26 patients with left bundle branch block, each using an emulator trained without that patient, and the calibrated posteriors were compared against the full simulator.

**Results:** A single emulator, trained without the held-out patient, calibrated each patient’s digital twin in 59.7–63.0 s per patient, versus 15 h with the simulator. Resampling the real cohort was the more efficient training strategy, bringing every output to *R*^2^ *≥* 0 .9, which augmentation did not achieve for the same simulation budget. For held-out patients, both strategies performed comparably (median leave-one-out normalized root-mean-square error 11.0% vs. 10.7%; *p*=0 .745), and error rose with distance from the cohort centroid (Spearman *ρ* = +0 .406). The calibrated models showed no cohort-wide bias (median standardized residual *−*0 .12between simulator and emulator), and 95% predictive intervals contained 78.5% of targets, with the largest misfit in systolic blood pressure.

**Conclusion:** Removing the need for new simulations for every new patient is a meaningful step toward uncertainty-aware cardiac digital twins that fit within a routine clinical workflow. Because the simulator, surrogate and sampler are interchangeable, the approach used here can be applied to other (cardiac) models and cohorts.

## 1 INTRODUCTION

Cardiovascular medicine is increasingly moving away from population-based guidelines towards individualized care tailored to each patient’s specific physiology^1^. Realizing this vision requires patient-specific cardiac models: multiscale, biophysical digital twins of the heart and circulation that integrate patient imaging, electrocardiography, and/or hemodynamic data to reproduce clinically measured biomarkers and simulate individual pathophysiology^1–6^. These models could guide therapies, for example by predicting a patient’s response to cardiac resynchronization therapy (CRT)^7^ or planning the ablation of ventricular tachycardia^8^, and enable in silico trials on virtual patient cohorts^9,10^. In an ideal scenario, a digital twin is to be generated or updated within minutes of new data becoming available, with an honest estimate of its uncertainty, so that it fits within a routine clinical workflow. Yet in practice, building each patient-specific model remains time- and resource-intensive, especially when including uncertainty quantification, which is essential for safe clinical use.

Uncertainty quantification is one of the main hurdles towards routine deployment of uncertainty-aware digital twins in the clinic. Bayesian calibration offers a mathematical framework for building patient-specific models, inferring parameters from clinical data while quantifying the uncertainty in those estimates by methods such as Markov chain Monte Carlo (MCMC) sampling or Bayesian history matching (BHM)^11–14^. Yet, these approaches require thousands of model evaluations per patient, far more than a clinical time frame allows when each evaluation requires a full biophysics model simulation. The payoff of the increased computational cost is that, in contrast to traditional deterministic methods, Bayesian methods yield probabilistic predictions with credible intervals that communicate confidence in model-derived biomarkers and therapy outcomes^9,15,16^, reflecting uncertainty that can arise from biological variability, sparse and noisy clinical measurements, and modeling assumptions across mechanics, electrophysiology, and long-term remodeling^17–22^.

Much of the cost of Bayesian calibration of cardiac digital twins can be removed by making each model evaluation cheaper. Surrogate models do so by replacing the simulator with a statistical emulator, such as a Gaussian process or neural network, trained on a limited set of simulations. They accelerate uncertainty quantification and parameter inference by orders of magnitude while preserving accuracy for key clinical quantities of interest^2,18,23^, and can be paired with MCMC or Bayesian history matching to identify parameter posteriors. The simulator can be a high-fidelity model or a reduced-order model, which lowers the cost further by simplifying the underlying physics while retaining the essential mechanisms of cardiovascular function. Reduced-order models have been used extensively to study cardiovascular mechanisms^24–28^ and, increasingly, in precision-medicine approaches^7,29^, and a surrogate trained on one is cheap both to train and to evaluate.

However, a key limitation remains that surrogate models are typically trained on a patient-by-patient basis. This is because the emulator is trained to reproduce a model’s output that is a function of not just the free parameters, but also of patient-specific constants such as cardiac wall mass and local activation timing. These are measurable and thus fixed but still vary across individuals, so an emulator trained on one patient’s constants does not generalize to another’s. Therefore, generating a new patient-specific model still requires generating fresh training simulations for each calibration. This intensive computational cost currently stands between Bayesian model calibration and near-instant patient-specific model calibration in routine clinical practice.

The goal of this work is to personalize a cardiac model for a new patient at little or no additional simulation cost, by training a single emulator that exploits structure shared across a patient cohort. Unlike a per-patient emulator, this *population-level* emulator shares training across patients, so that calibrating a patient requires no new high-fidelity simulations at the moment of calibration. We hypothesized that a population-level emulator yields accurate patient-specific calibration with quantified uncertainty, and that it is more accurate for patients whose clinical features are close to the training cohort.

Specifically, we contribute (i) a population-level Gaussian process emulator (GPE) whose inputs include a low-dimensional latent encoding of each patient’s clinical features, so that one set of trained emulators serves the whole cohort; (ii) a procedure to determine the optimal training strategy of the emulator that generalizes beyond the combination of cohort, emulator, and simulator used in this study; (iii) an MCMC calibration pipeline that returns converged posteriors in about a minute per patient, including for patients the emulator was not trained on; and (iv) validation of the calibrated posteriors against the full biophysical model. We demonstrate the framework on a 26-patient CRT cohort and release it as an update to our open-source Python calibration package CHAMELEON^14^.

## 2 METHODS

### 2.1 Clinical data

Data from 26 patients with left bundle branch block (LBBB) were used, coming from three clinical studies at the University of Virginia. All participants had chronic systolic heart failure, a left ventricular ejection fraction *≤* 35%, and a guideline class I or II indication for CRT. Informed consent was obtained from all patients, and the studies were approved by the University of Virginia Institutional Review Board for Human Subjects Research; full inclusion and exclusion criteria are given in^30,31^.

Prior to CRT, each patient underwent non-invasive measurements that we divide into two sets according to how they enter the model. The first,

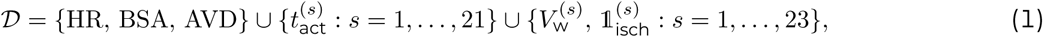

collects measurements with direct counterparts in the model that could thus be prescribed directly (Appendix Table A1). Heart rate (HR) was calculated from a 12-lead ECG and body surface area (BSA) from the Du Bois formula. The remaining quantities were obtained per functional segment, dividing the heart into 23 segments: one each for the right atrium (RA) and left atrium (LA), five for the right free wall (RFW), and, following the 17-segment AHA model of the left ventricle without the apical cap (segment 17)^32^, 11 for the left free wall (LFW) and five for the septal wall (SW); segments *s* = 1 *, … ,* 21 are ventricular and *s* = 22 , 23 atrial. For each segment *s* we determined the wall volume 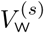 from cine MRI; a binary ischemia indicator 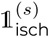 (true if *≥* 50% of the segment was ischemic) from late gadolinium enhancement (LGE) MRI; and for each ventricular segment the activation time 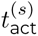 from displacement encoding with stimulated echoes (DENSE) MRI, the five RFW segments were assigned the same activation time in every patient (Appendix Table A1). The atrioventricular delay (AVD) was defined as the interval between atrial activation and the earliest ventricular activation. HR is prescribed to the simulator only. The other measurements form the patient feature vector 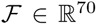 used by the emulator (Section 2.3.1): BSA and, for each of the 23 segments, the activation time, wall volume and ischemia indicator. Activation times in *F* are expressed relative to the earliest ventricular activation, so the two atrial entries carry the AVD.

The second set,

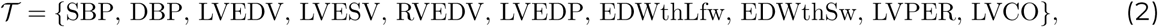

collects measurements without direct model counterparts, used as calibration targets (Section 2.4; Table 1). Cuff measurements provided systolic (SBP) and diastolic (DBP) blood pressure; cine MRI provided left ventricular (LV) end-diastolic (LVEDV) and end-systolic volume (LVESV), right ventricular end-diastolic volume (RVEDV), end-diastolic left free wall (EDWthLfw) and septal wall end-diastolic wall thickness (EDWthSw), LV peak ejection rate (LVPER), and cardiac output (LVCO). LVCO is the volumetric cardiac output reported by the analysis software (SuiteHeart), i.e. the product of stroke volume (LVEDV *−* LVESV) and heart rate. Therefore, LVCO is not independent of the volumetric targets: stroke volume contributes to the likelihood through both, which up-weights it relative to the other targets. We retain LVCO to reflect the clinical measurement set. MONARCH, in contrast, computes LVCO from the net forward flow through the aortic valve. At the posterior draws, this flow-based value differed from MONARCH’s own volumetric value by a median of 4.0%, but by more than 10% for about a quarter of draws (90th percentile 14.8%). Additionally, LVEDP was not measured but estimated from NYHA functional class via pulmonary capillary wedge pressure (Appendix A2); we included it to provide a target that constrains the pressure at end diastole, which the volumetric targets alone leave undetermined. The uncertainties in Table 1 are one standard deviation of the measurement error. For the cuff pressures we used 8 mmHg, the largest standard deviation of the device–reference difference that the universal validation standard for blood pressure devices accepts^33^. Inter-observer differences in CMR left ventricular volumes are typically 5 to 11 mL^34–36^, so we used 10 mL for LVEDV and LVESV; right ventricular contours are traced less consistently than left ventricular ones^36^, so we used 20 mL for RVEDV. Observers place the left ventricular myocardial contours about 1.2 mm apart on average^36^, which motivates the 1 mm uncertainty of the wall thicknesses. The uncertainty of LVEDP is the spread of pulmonary capillary wedge pressure within a NYHA class (Appendix A2). For LVPER and LVCO, for which we found no reported measurement error, we used roughly 10% of the cohort median.

**Table 1:** Clinical measurements comprising set *T*, used as targets for the parameter calibration, given as median [min, max] over the 26 patients. The sources of the uncertainties are given in Section 2.1.

| Target | Description | Value | Uncertainty<br>(1 SD) | Units |
|---|---|---|---|---|
| SBP | Systolic blood pressure | 114 [78, 150] | 8.0 | mmHg |
| DBP | Diastolic blood pressure | 68 [34, 100] | 8.0 | mmHg |
| LVEDV | Left ventricular end-diastolic volume | 234 [100, 406] | 10 | mL |
| LVESV | Left ventricular end-systolic volume | 178 [75, 366] | 10 | mL |
| RVEDV | Right ventricular end-diastolic volume | 125 [73, 250] | 20 | mL |
| LVEDP | Left ventricular end-diastolic pressure | 15 [15, 23] | 5.0 | mmHg |
| EDWthLfw | Ventricular left free wall end-diastolic wall thickness | 7.77 [4.65, 12.08] | 1.0 | mm |
| EDWthSw | Ventricular septal wall end-diastolic wall thickness | 8.12 [5.28, 11.06] | 1.0 | mm |
| LVPER | Left ventricular peak ejection rate | 500 [38, 1069] | 50 | mL/s |
| LVCO | Left ventricular cardiac output | 3.87 [0.54, 6.66] | 0.50 | L/min |

### 2.2 Biophysics model

As a testbed for the framework, we used MONARCH, our open-source Python implementation of a reduced-order model of the heart and circulation that was previously used to simulate LBBB, CRT, atrial fibrillation, and mitral valve regurgitation^14,25,37^. In this model, the four chambers are represented by thick-walled spherical segments following the TriSeg method^38^: the ventricles are modeled as three spherical segments (LFW, RFW, and SW) coupled through tension equilibrium at a shared junction, and the LA and RA as single spheres, mechanically uncoupled from the ventricles. Each wall segment uses a Hill-type model that generates an active myofiber stress scaled by a contractility parameter^38,39^ and a passive elastic stress that depends on fiber stretch^25^. Each ventricular wall is functionally divided into multiple segments using the MultiPatch method^40^, allowing for dyssynchronous activation and mechanical dyssynchrony that is a hallmark of LBBB. The systemic and pulmonary circulations are each modeled as a five-element RCRCR Windkessels, with pressure-sensitive valves enforcing one-way flow and a stressed blood volume setting the operating pressures. The coupled equations of heart and circulation are integrated in a closed loop system, yielding chamber pressures, volumes, and wall mechanics over the cardiac cycle. We selected the free parameters by Morris elementary-effects screening^41^ of the 12 candidate model parameters against the outputs in *T*. The mean absolute elementary effect *µ^∗^* of each parameter was normalized by the range of each output and averaged across outputs, and we retained the smallest set of parameters whose cumulative normalized effect reached 90% of the total. This yielded seven free parameters (Table 2):

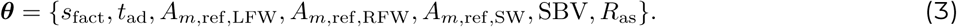

**Table 2:** Biophysics model parameters retained for calibration after Morris screening. These seven parameters constitute the free parameter vector ***θ***. Prior range, bounds of the uniform calibration priors; training range, the hyperrectangle within which training parameters were drawn (Section 2.3.2).

| Parameter | Description | Prior range | Training range | Units |
|---|---|---|---|---|
| *s*<sub>fact</sub> | Active stress coefficient | 0.025–0.350 | 0.045–0.349 | MPa |
| *t*<sub>ad</sub> | Active contraction duration | 50–450 | 50.2–444.1 | ms |
| *A*<sub>*m*,ref,LFW</sub> | LFW midwall reference area | 6,000–20,000 | 6,342–16,645 | mm<sup>2</sup> |
| *A*<sub>*m*,ref,RFW</sub> | RFW midwall reference area | 8,000–20,000 | 8,336–19,546 | mm<sup>2</sup> |
| *A*<sub>*m*,ref,SW</sub> | SW midwall reference area | 2,000–10,000 | 2,784–8,991 | mm<sup>2</sup> |
| SBV | Stressed blood volume | 1,000–3,000 | 1,005–2,951 | mL |
| *R*<sub>as</sub> | Systemic arterial resistance | 0.1–3.0 | 0.106–2.989 | mmHg s mL<sup>−1</sup> |

The remaining parameters were set constants *C* based on previous studies (Table A2).

### 2.3 Population-level emulator

Surrogate models like GPEs can reduce the computational cost of each model evaluation by several orders of magnitude^42^, but are conventionally trained on one patient at a time^43–45:^ because the emulator response depends not only on free model parameters ***θ*** (Section 2.2) but also on patient features *F* (Section 2.1), each patient nominally needs its own emulator, so the training cost grows linearly with the number of patients. We here describe a framework to train a single cohort-level emulator that takes a principal component analysis (PCA) encoding of *F* as additional input, so that one trained emulator serves the entire cohort and can be applied to new patients without retraining.

#### 2.3.1 Latent representation of patient features

Across a clinical cohort, patient features are far from independent of one another; they reflect shared anatomical and physiological structure, such as the coupling between chamber geometry and body surface area or the correlation of regional activation delays. Using all features directly as emulator inputs therefore creates a higher-dimensional input space than necessary, which complicates and slows emulator training. In our clinical application the feature vector of each patient, 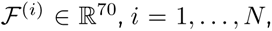, holds the BSA and the activation time, wall volume and ischemia indicator of each of the 23 segments (Section 2.1), and is subsequently reduced by PCA. Each feature is first standardized to zero mean and unit variance across the cohort, placing features measured in different units on a common scale. Features that are constant across the cohort, such as an ischemia indicator 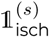 that is zero for every patient, have zero variance and carry no information to distinguish patients, and are thus eliminated entirely by the dimensionality reduction. In our cohort, 16 of the 70 features were constant: the five RFW activation times and the ischemia indicators of the 11 segments that were never ischemic (Appendix Table A1). We then decompose the matrix of standardized features **X** = **UΣV***^⊤^* and retain the *k* leading components that together explain at least 95% of the variance. Each patient is then summarized by the *k*-dimensional projection

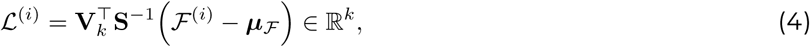

with ***µ****_F_* the cohort mean and **S** the diagonal matrix of cohort standard deviations (set to one for constant features). In our CRT cohort (*N* = 26), this criterion yields *k* = 12, i.e. *F* ∈ ℝ^70^ → *L* ∈ ℝ^12^. The emulator is therefore trained on the joint input (***θ****, L*) *∈* ℝ^19^ rather than (***θ****, F*) *∈* ℝ^77^, a far more tractable learning problem.

#### 2.3.2 Emulator training

Training data are generated by running the simulator once per (patient, parameter) pair. For each patient *i*, free parameters ***θ*** are drawn by Latin hypercube sampling within the training ranges of Table 2, the hyperrectangle spanned by the 1st to 99th percentiles of the parameter posteriors from an initial calibration of every patient, pooled over the cohort, which lies inside the prior ranges, and MONARCH is evaluated for the joint input (***θ****, D*^(*i*)^*, C*). These simulations dominate the computational cost of the framework, but only need to be paid once, upfront. Each evaluation yields the simulated outputs **y**^(*i*)^ *∈* ℝ^10^, the model counterpart of the ten calibration targets in *T* (Section 2.1). Simulations that produce outputs outside physiological bounds or fail to converge (Supplemental Material, Table S1) are discarded.

While the simulator requires the patient’s measurements *D*, the emulator uses the low-dimensional latent encoding *L* of the features *F*. Each simulated output is therefore paired with (***θ****_j_, L*^(*i*)^) to form up to *n_s_* training examples *{*(***θ****, L*^(*i*)^, **y**^(*i*)^)*}* per patient or chimera. We train a separate GPE for each of the ten outputs as a function of this joint input (***θ****, L*),

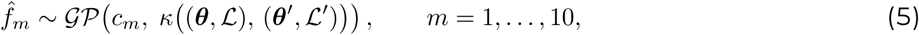

following the standard practice of emulating each output independently^14,46,47^. Inputs and outputs are normalized using z-scores so the constant mean function is effectively *c_m_ ≈* 0. The covariance kernel *κ* and its hyperparameters per output are selected by cross-validated kernel comparison using AutoEmulate^48^, and the hyperparameters are fitted by maximizing the marginal likelihood with the Adam optimizer for up to 200 epochs (100 for the emulators trained with augmentation in the leave-one-out comparison in Section 3.2), stopping early once the relative improvement in the loss has stayed below 10*^−^*^4^ for 20 consecutive epochs; the framework is agnostic to this choice, and the optimal kernel will generally differ across models, datasets, and input and output dimensionality. At inference the Gaussian process (GP) posterior for each output is Gaussian with closed-form mean and variance, providing both a prediction and its uncertainty^49^. Surrogate models other than GPs can also be used, however the GP posterior mean is analytically differentiable with respect to its inputs, so this gradient is available in closed form and gradient-based samplers can be used for calibration.

#### 2.3.3 Training design: resampling and augmentation

The computational cost *C*, the number of simulations run to train the emulator, can be spent in two ways: by sampling the real patients more densely in parameter space, or by covering more of the latent space with synthetic patients. We compare two strategies that start from the same baseline of one simulation per real patient (*C* = *N*). The first strategy, *resampling*, keeps the *N* real patients and draws *n_s_* parameter vectors ***θ*** per patient using Latin hypercube sampling, so that *C* = *N · n_s_*. The second strategy, *augmentation* keeps one parameter draw per real patient and spends every additional simulation on a new synthetic patient, with a single parameter draw, so that *C* = *N* + *Ñ*, where *Ñ* is the number of synthetic patients. Resampling adds information only about the real patients, while augmentation adds a new point in latent space. Resampling adds information only about the real patients, whereas each synthetic patient adds a new point in latent space. With *N* = 26 patients spread over *k* = 12 latent dimensions, the coverage of the cohort in latent space is sparse, so augmentation tests whether denser coverage improves the emulator’s accuracy for new patients.

We call the synthetic patients digital chimeras, after Dou et al.^50^ coined the term for virtual cardiac anatomies assembled from the anatomical substructures of different subjects. Each of our chimeras is instead the midpoint of the complete feature vectors of two members *i* and *j* of the current pool, which holds the real patients and the chimeras selected so far,

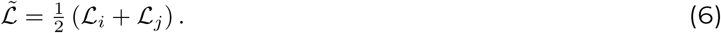

Because the PCA projection (Eq. 4) is affine, this corresponds to averaging the parents’ features elementwise, which gives the chimera’s simulator inputs; binary ischemia indicators are kept binary rather than averaged: where the two parents differ in a segment, both values are generated as separate candidates and the selection below chooses between them (Supplemental Material). Interpolating between existing samples to create synthetic training data is an established technique in machine learning^51^. We selected chimeras by greedy maximin, or farthest-point, selection^52,53^: at each step, we accepted the candidate farthest from its nearest real patient or previously accepted chimera. The candidates are initially the 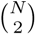 midpoints between pairs of real patients, and each acceptance adds the midpoints between the new chimera and every existing pool member. Coverage therefore keeps improving as chimeras are added, rather than plateauing once the pairwise midpoints are used up. Apart from its ischemia indicators, every chimera is a convex combination of real patients, so augmentation adds coverage within the cohort without extending it (Fig. 1), and a new patient outside the convex hull of the real patients remains an extrapolation for the emulator. The Supplemental Material describes how the simulator inputs are constructed, how ischemia is handled, and the pruning that keeps the growing candidate pool tractable.

**Fig. 1:**
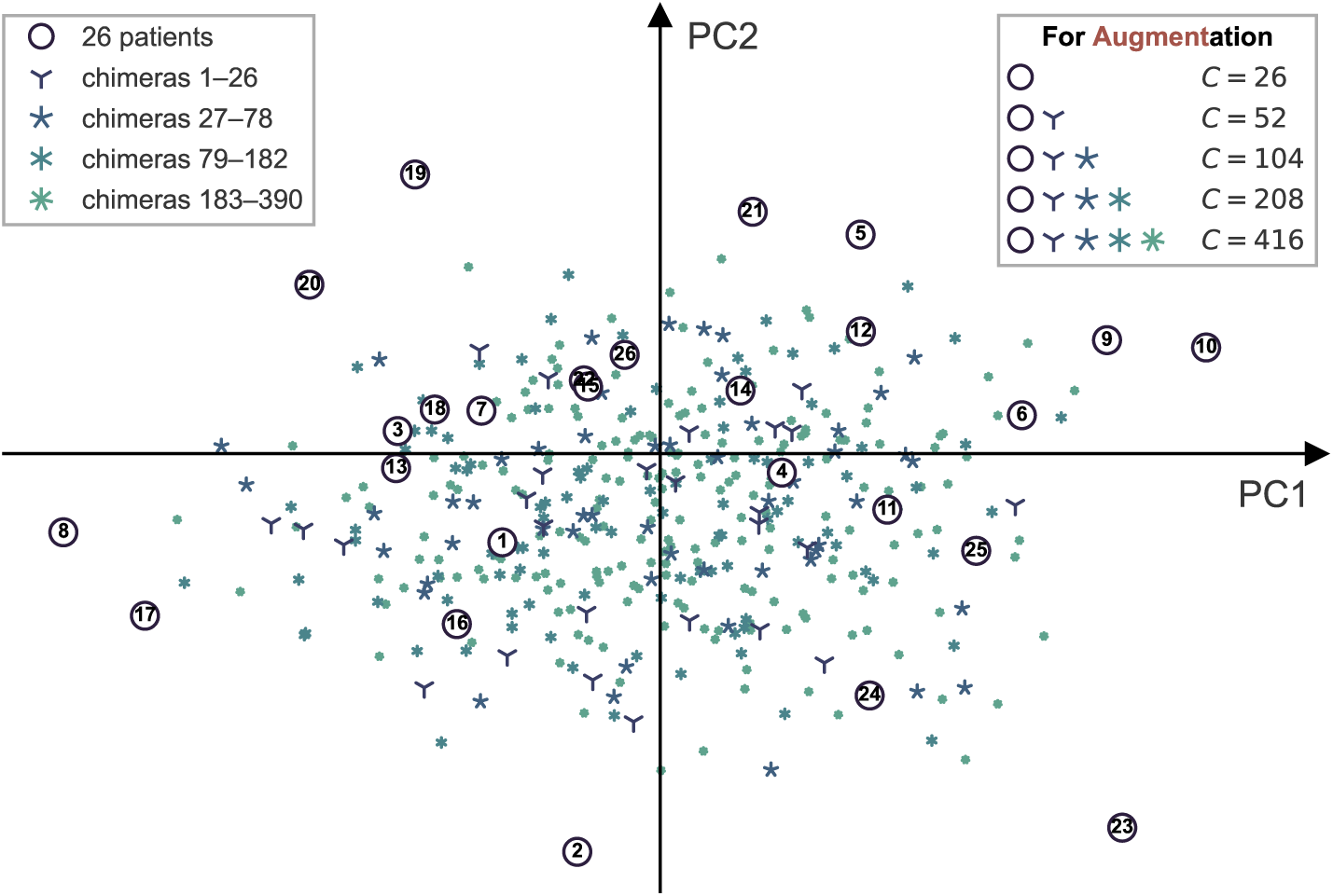
Digital chimeras fill under-sampled regions of the latent patient-feature space. Real patients (numbered circles) and chimeras on the first two principal components (PC1, 37.6%; PC2, 12.0% of feature variance). Chimeras are shown by the batch in which they were added as the cost of the Augment strategy doubled from *C* = 52 to 416 (*Ñ* = 26 to 390). No chimera extends the region spanned by the real patients. The calibration emulators use no chimeras.

### 2.4 Calibration

The goal of the calibration is to infer for each patient *i* the values of the free parameters ***θ***that make the model reproduce the clinical targets *T* ^(*i*)^. We employ a Bayesian approach and infer the joint posterior *p*(***θ*** *| T* ^(*i*)^), which is proportional to the product of two terms: the *likelihood p*(*T* ^(*i*)^ *| **θ***), measuring how well a given ***θ*** reproduces the measured targets, and the *prior p*(***θ***), encoding what is known about plausible parameter values beforehand. We here chose uniform priors over the bounds in Table 2 to treat all parameter combinations as equally likely a priori. Estimating the joint posterior rather than a single best fit is what allows the calibrated model to account for measurement uncertainty, which is crucial for providing realistic and trustworthy clinical predictions.

We treat each target 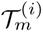 as a noisy observation of the corresponding emulator prediction,

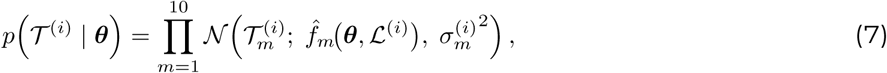

where *N* (*x*; *µ, σ*^2^) denotes the normal probability density of *x* with mean *µ* and variance *σ*^2^, and 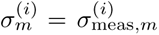 is the clinical measurement uncertainty of target *m* (Table 1), which does not depend on ***θ***. Up to an additive constant, the log-likelihood is therefore a weighted sum of squared residuals,

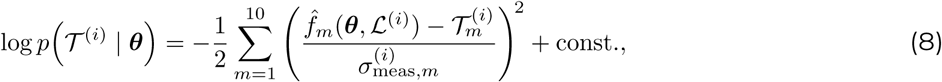

in which targets with the lowest uncertainty weigh more. The emulator’s own predictive uncertainty *σ̂*_GP_*_,m_*(***θ****, L*^(*i*)^) is not included: the emulator mean is treated as exact during sampling, and emulator error is assessed after calibration by evaluating the posterior draws with MONARCH (Section 2.5). Bayesian calibration differs from deterministic least-squares fitting in returning the full set of parameter values consistent with the data rather than a single best fit.

Because the posterior does not have a closed form and we cannot sample it directly, we explore it with Markov chain Monte Carlo (MCMC): a method that walks through parameter space, preferentially visiting parameter sets that fit the data well, and collects the visited sets as samples. The resulting collection approximates the posterior—the more often a region is visited, the more probable it is. Because we use an emulator instead of a simulator, the time to evaluate each likelihood drops to mere milliseconds, making full MCMC sampling per patient feasible. We sampled the posterior with the differential-evolution Metropolis sampler differential evolution Metropolis variant (DEMetropolisZ)^54^, which runs several interacting chains, proposes moves from differences between past states of the chains, and needs no gradients. Like the choice of emulator, the choice of sampler is not intrinsic to the framework and depends on factors such as the dimensionality of ***θ*** and whether the emulator is differentiable. Because the GP posterior mean is differentiable, gradient-based samplers such as the No-U-Turn Sampler (NUTS)^55^ can be used instead. We assessed convergence with the rank-normalized split-*R̂* and the bulk effective sample size^56,57^, computed over all chains, and compared them with the recommended thresholds of *R̂ ≤* 1.01 and a bulk effective sample size of at least 400^57^; we also report the tail effective sample size. We used 4 chains of 20,000 draws after 10,000 tuning steps on a single CPU core. The tuning draws of served as burn-in and were discarded; all remaining draws were used. Diagnostics for every patient and trace plots for three representative patients are given in the Supplemental Material, Section S3.

### 2.5 Emulator assessment

To identify the optimal configuration of the emulator in terms of accuracy and computational efficiency, we assessed the impact of the number of samples of ***θ*** per patient *n_s_* and the number of chimeras *Ñ* (Section 2.3.3). At each setting we trained a single population emulator on all *N* = 26 patients (together with the *Ñ* chimeras) and evaluated its accuracy on held-out parameter samples: for every patient we reserved a fixed set of 64 of its parameter samples—the same held-out set for all costs and for both the resampling and augmentation strategies—and trained only on the remaining samples, so that no graded sample was ever used in training. These samples came from a separate pool of 256 Latin hypercube samples per patient; simulations that failed or fell outside physiological bounds were dropped rather than redrawn, leaving between 21 and 58 valid held-out samples per patient (median 43). Accuracy was averaged as the median over repeated draws of the training samples, with the heldout set fixed. The computational cost is the number of simulator evaluations, *C* = (*N* + *Ñ*) *n_s_*, counted before invalid simulations were dropped. Emulator accuracy was assessed by calculating *R*^2^ and the normalized root-mean-square error (NRMSE) between emulator and simulator, defined as

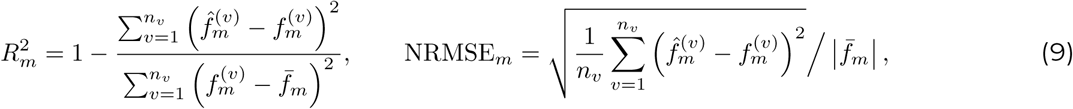

with 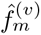 and 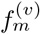 the emulator and simulator outputs *m* for the *v*-th held-out sample, *n* the number of held-out samples, and *f̅_m_* the mean of 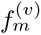 over those samples. Overall accuracy per measurement andper patient was assessed by calculating the median and interquartile range of *R*^2^ and NRMSE*_m_*. The two configurations (resampling and augmentation) were compared per patient with a paired Wilcoxon signed-rank test, and the influence of a patient’s position in the cohort on emulator accuracy was quantified by Spearman’s rank correlation with its latent distance to the cohort mean and its mean latent distance to the three nearest real patients.

Calibration accuracy was assessed by how well calibrated patient-specific models reproduce the clinical data. To test the framework’s ability to generate personalized models for unseen patients, each patient was calibrated with the leave-one-out cross-validation (LOOCV) emulator that excluded that patient. For each patient, we drew 100 parameter sets from the posterior and simulated them with MONARCH, using the patient’s measurements *D*^(*i*)^, and compared these simulations with the emulator predictions for the same draws. Emulator fidelity was measured by *R*^2^ and NRMSE between emulator and MONARCH outputs. Agreement with the clinical data was measured by the share of targets within the 95% posterior predictive interval, computed from the MONARCH outputs at the posterior draws with measurement noise added 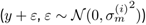, and by the standardized residual,

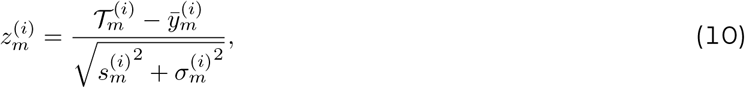

with 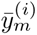 and 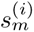 the mean and standard deviation of the posterior predictive MONARCH outputs and 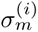 the clinical measurement uncertainty of target *m* (Table 1). The denominator thus combines parameter and measurement uncertainty, so that *|z| <* 2 indicates a target consistent with the calibrated model given both. Finally, the posterior contraction 1 *− σ*_post_*/σ*_prior_ quantified how strongly the targets constrain each parameter (Supplemental Material, Section S4), with *σ*_prior_ and *σ*_post_ the standard deviations of the uniform prior and the posterior.

### 2.6 Implementation

All methods were implemented in Python and will be made publicly accessible through our lab’s GitHub page upon publication of this manuscript. We include documentation and curated Python files and Jupyter notebooks to reproduce all results. The MONARCH package was used for the cardiac simulator and CHAMELEON for emulation and calibration. These packages use PyMC for MCMC sampling; NumPy and Pandas for calculations, data processing and analysis; MatPlotLib, Seaborn, and ArviZ for visualization; and AutoEmulate, PyTorch, and NumPy for GP emulator training and evaluation. For the latter, we exploited the analytical form of the GP posterior to implement a NumPy alternative for the trained PyTorch implementation, resulting in a *∼*200-fold acceleration. The one-time generation of training simulations, the cross-validation experiments (Sections 3.1 and 3.2), the MONARCH posterior predictive checks were run in parallel on HPC3, the high-performance computing cluster of the University of California, Irvine. Calibrating an unseen patient with a trained emulator does not require a cluster: each held-out calibration took about a minute on one CPU core of a laptop (Apple M1 Max), the machine on which the MONARCH simulation time (0.44 s) was also measured. A large language model (Claude Opus 5 and 5.5, Sonnet 5 and 5.5, Anthropic) was used to generate and debug analysis code, verified by the authors.

## 3 RESULTS

Having defined the framework in Section 2, we validate it on the clinical cohort through a sequence of experiments, each addressing one design decision and motivating the next. Although we test the framework on a specific cohort and simulator, the procedure used to configure and validate it applies to any combination of the two. We first ask how a fixed computational cost is best spent to train the emulator, on sampling real patients more densely or on adding digital chimeras, and how many simulations are needed to reach acceptable accuracy (Section 3.1). We then examine, patient by patient, how accurately the resulting emulator predicts patients it was not trained on, and what governs that accuracy (Section 3.2). Finally, we use the optimized emulator to calibrate the model for every patient and verify the personalized models against the full simulator (Section 3.3).

### 3.1 What is the optimal training strategy for the population-level emulator?

First, we asked how a computational cost *C*, the number of simulations, is best spent: on more simulations of the real patients (Resample) or on digital chimeras (Augment; Section 2.3.3). Both strategies start from one simulation per real patient (*C* = 26), and at every cost both were scored (Section 2.5) on the same held-out parameter samples of every patient.

Both strategies improved quickly at first, with diminishing returns as the cost grew (Fig. 2). Resample was the more efficient strategy: it brought every output to *R*^2^ *≥* 0.9 at *C* = 832, whereas Augment had not done so at the highest cost tested, where its worst-predicted output (DBP) reached *R*^2^ = 0.836 (Fig. 2a). We did not test higher costs, so Augment might reach this level with a further doubling; alternatively, its gains may level off once the chimeras saturate the interior of the latent space, where additional chimeras add little new information (Section 4.3). From *C* = 104 onward, Resample also had the lower median per-output NRMSE at every cost level (Fig. 2b). At *C* = 3,328, per-output *R*^2^ ranged from 0.956 (LVCO) to 0.999 (EDWthLfw) under Resample, and the median per-output NRMSE was 5.1% (IQR 4.2–6.3%), compared with 11.9% under Augment (Fig. 2c, d).

**Fig. 2:**
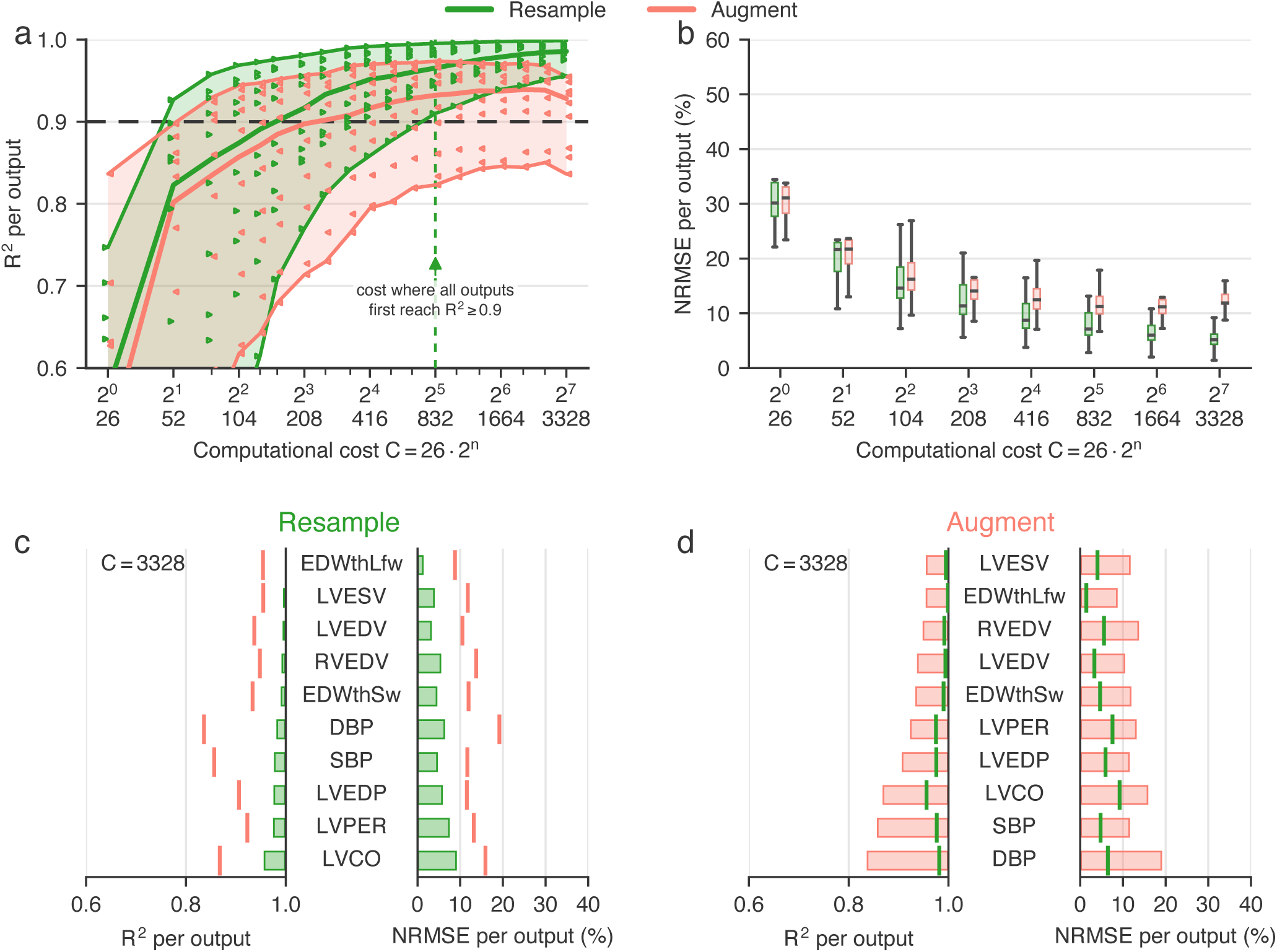
Resampling the real cohort reaches uniform accuracy at lower cost than chimera augmentation. Emulator accuracy against computational cost *C* when the additional simulations are spent on more parameter samples of the 26 real patients (Resample) or on chimeras (Augment), scored on 64 held-out parameter samples per patient. **(a)** *R*^2^ of the ten outputs; bands span the best- to worst-predicted output and thick lines show the median; values below 0.6 are not shown. The vertical line marks the cost at which every output first reaches *R*^2^ = 0 .9 under Resample. **(b)** NRMSE of the ten outputs. **(c, d)** Per-output *R*^2^ and NRMSE at *C* = 3,328 for Resample (c), the configuration used for calibration, and Augment (d); tick marks show the other strategy. NRMSE, normalized root-mean-square error.

For the training cohort, resampling the real patients thus reached a given accuracy at lower cost than augmentation. Whether chimeras help for patients outside the training set is tested in Section 3.2. We therefore used the Resample strategy for the calibration emulators, with *n_s_* = 128 parameter draws per real patient and *Ñ* = 0 (*C* = 26 *·* 2^7^ = 3,328 for the full cohort; 3,200 simulations per LOOCV fold). This is two doublings beyond the cost at which every output first reached *R*^2^ *≥* 0.9 under Resample (*C* = 832 = 26 *·* 2^5^; Fig. 2a), leaving a margin for patients the emulator was not trained on, for whom accuracy is lower (Section 3.2).

### 3.2 Does the emulator generalize to new patients, and what governs its accuracy?

Because the central advantage of a population-level emulator is that it can be applied to a new patient without retraining, we next tested how well it predicts patients excluded from training. In leave-one-out cross-validation (LOOCV), each patient was held out in turn and predicted by the emulator trained on the remaining 25 real patients. We compared the two training strategies at the same computational cost (3,200 simulations per fold): resampling (*n_s_* = 128 per patient, *Ñ* = 0) and augmentation (*n_s_* = 1 per patient and *Ñ* = 3,175 chimeras with one simulation each, regenerated from the remaining 25 patients only; Section 2.5).With resampling, the median LOOCV NRMSE across patients and outputs was 11.0% (IQR 6.8–15.7%), with per-output medians from 6.5% (LVESV) to 21.3% (DBP). Augmentation performed comparably, with a median of 10.7% (IQR 7.3–15.6%; Fig. 3b). Augmentation reduced the NRMSE for 12 of 26 patients, and the median paired difference (augmentation minus resampling) was +0.83 percentage points (Wilcoxon signed-rank *p*=0.745). At equal computational cost, the two strategies were therefore equally accurate for held-out patients, whereas resampling was the more accurate strategy for the training cohort (Section 3.1).

**Fig. 3:**
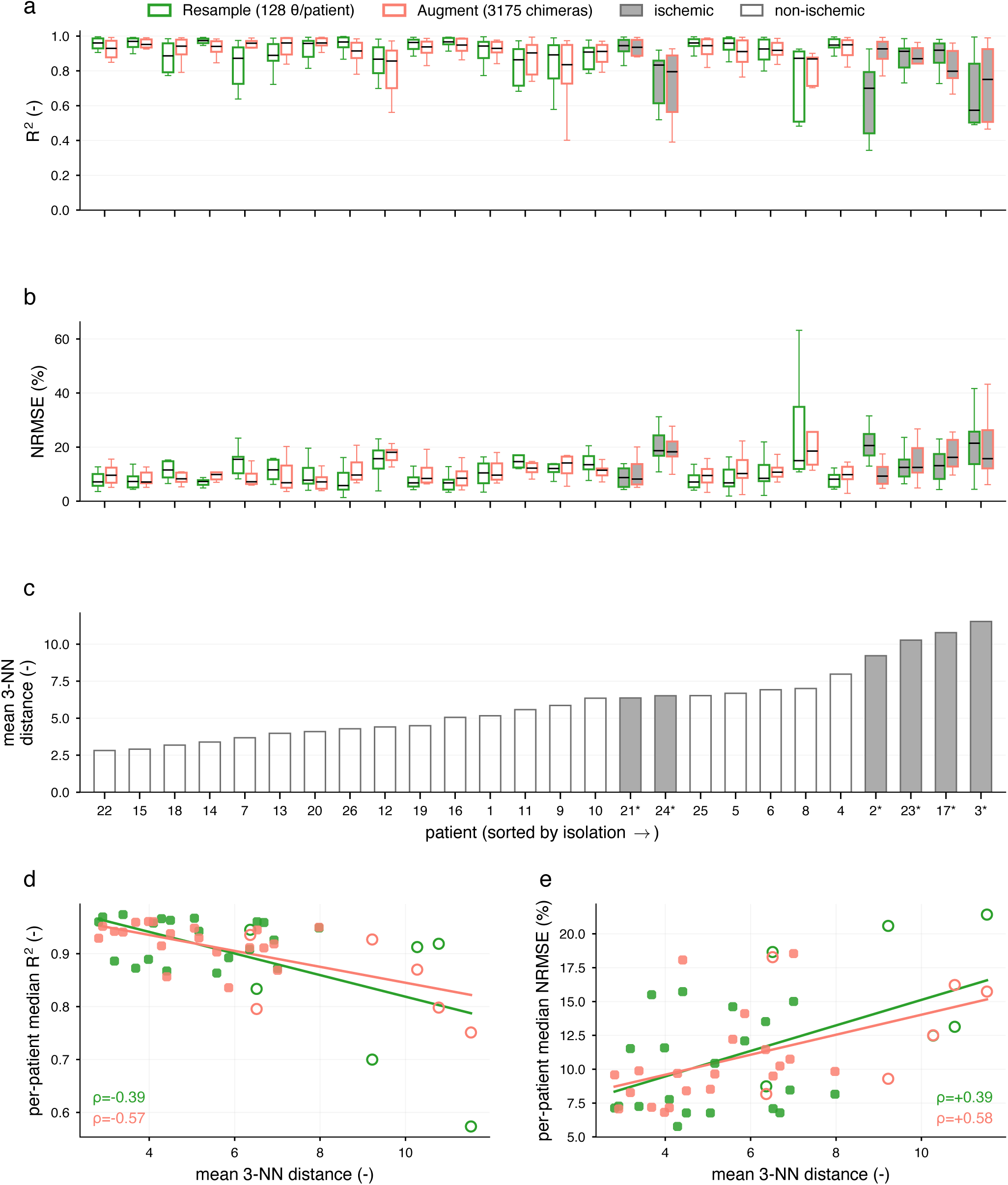
Resampling and augmentation give similar accuracy for held-out patients, and accuracy decreases with isolation in latent space. Each patient was predicted by Resample and Augment emulators trained on the other 25 patients at equal cost (3,200 simulations). Patients are ordered by their mean latent distance to the three nearest real patients. **(a, b)** *R*^2^ and NRMSE of the ten outputs. **(c)** Mean distance to the three nearest real patients. **(d, e)** Per-patient median *R*^2^ and NRMSE against that distance; *ρ*, Spearman’s rank correlation. Shading, outlines, red bars and asterisks mark the six patients with ischemic segments. NRMSE, normalized root-mean-square error.

Accuracy depended on a patient’s position in latent space (Fig. 3c–e). With resampling, per-patient NRMSE increased with distance to the cohort centroid (Spearman *ρ* = +0.406, *p*=0.040) and with mean distance to the three nearest real patients (*ρ* = +0.386, *p*=0.051), and per-patient median *R*^2^ decreased with that distance (*ρ* = *−*0.386, *p*=0.051). Patients far from the training cohort therefore tended to be predicted less accurately, as expected when a Gaussian process extrapolates. The four most isolated patients all had ischemic segments (Fig. 3c): because only 6 of the 26 patients had ischemic segments, a standardized ischemia indicator takes values near 5 for those patients, placing them far from the rest of the cohort in latent space.

Contrary to our hypothesis, chimera augmentation did not weaken this dependence; if anything, it was stronger with augmentation (Fig. 3d,e; centroid distance *ρ* = +0.529, *p*=0.005; three nearest patients *ρ* = +0.582, *p*=0.002; median *R*^2^ *ρ* = *−*0.571, *p*=0.002). For the most isolated patients the effect of augmentation was mixed: it lowered the median NRMSE of patient 2 from 20.6% to 9.3% but raised that of patient 17 from 13.1% to 16.2% (Fig. 3b).

### 3.3 Can the emulation framework personalize the model to clinical data?

Finally, we tested whether the population-level emulator can personalize the model to clinical data for patients it was not trained on. Each patient was calibrated with the emulator of the LOOCV fold that excluded that patient, trained with the resampling strategy (*n_s_* = 128, 3,200 simulations; Section 3.2), so neither the patient’s features nor its simulations were used to train the emulator that calibrated it. We used resampling because it was more accurate than augmentation for the training cohort (Section 3.1) and as accurate for held-out patients (Section 3.2). We then simulated 100 posterior draws per patient with the full MONARCH simulator.

Calibration took 59.7–63.0 s per patient (median 61 s) on a single CPU core. Of the 26 calibrations, 21 met the convergence criteria of *R̂ ≤* 1.01 and a bulk effective sample size of at least 400^57^, and the remaining five were only slightly outside them (*R̂ ≤* 1.020, bulk effective sample size *≥* 372; Supplemental Material, Table S2 and Figs. S1–S3).

The calibrated twins reproduced the clinical data for most patient–target pairs (Fig. 4). Simulated with MONARCH, 80.0% of pairs had *|z| <* 2, and the 95% posterior predictive interval, including measurement noise, contained the target for 78.5% of pairs. The volumetric targets were reproduced closely (median *z* = +0.09 for LVEDV and *−*0.05 for RVEDV), whereas SBP was overpredicted in many patients (median *z* = *−*2.42), which mostly reflects the reduced-order model rather than the emulator (Section 4.4). The poorest fit was for patient 11, for whom six of the ten targets had *|z| >* 2: the model could not reproduce this patient’s combination of targets jointly, most visibly underpredicting DBP (Fig. 4).

**Fig. 4:**
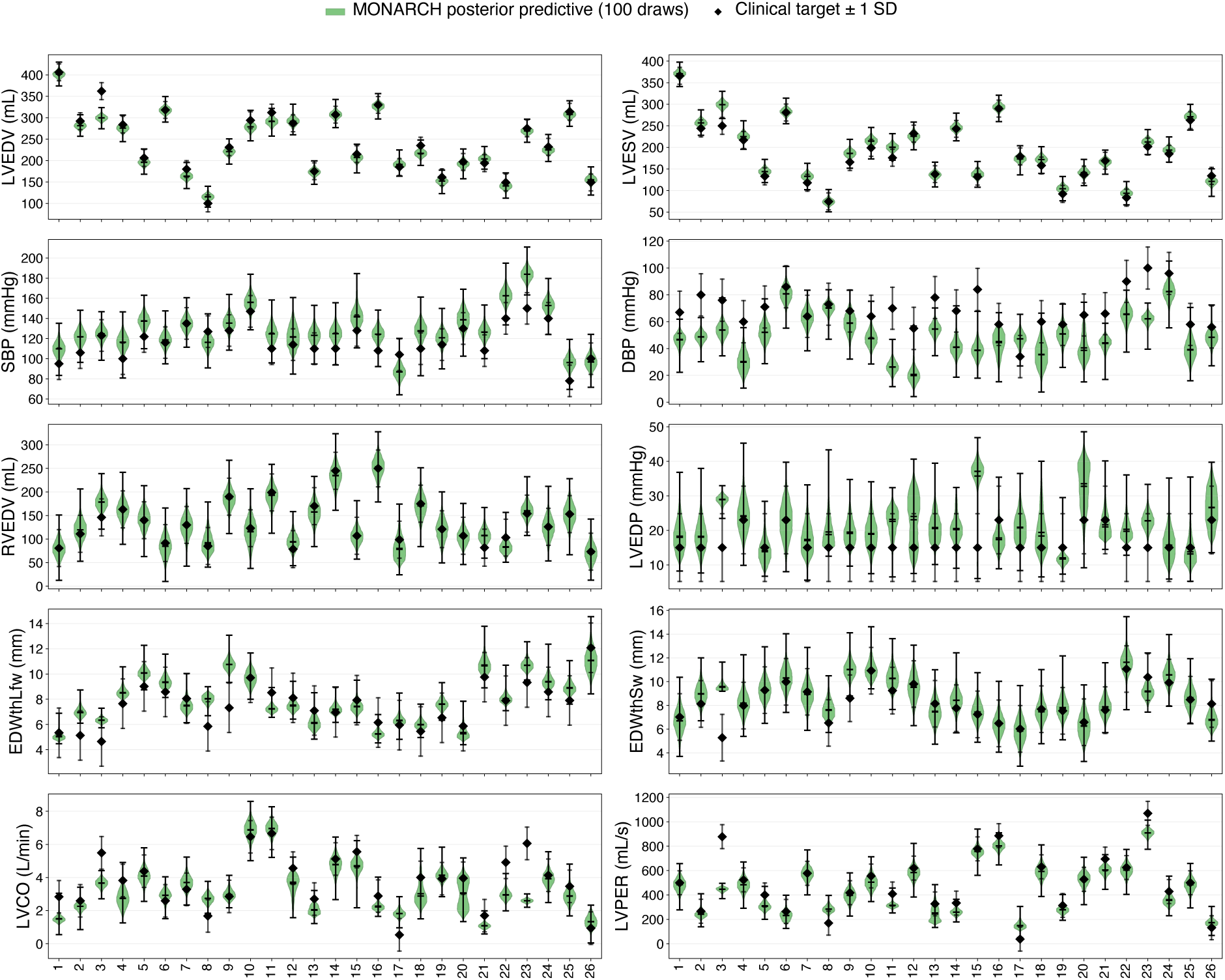
Posterior predictive distributions of the calibrated digital twins for held-out patients. MONARCH outputs at 100 posterior draws per patient (violins, without measurement noise) and the clinical targets *±* measurement uncertainty (diamonds). Each patient was calibrated with the leave-one-out emulator that excluded it. Abbreviations and units as in Table 1.

The emulator remained close to MONARCH in the posterior region (Fig. 5). At the posterior draws, the median NRMSE between emulator and MONARCH was 14.3%, and for nine of the ten outputs the mean emulator–MONARCH difference lay between *−*8.0% and +8.3% of the target; only SBP showed a larger bias (*−*22.5%; Fig. 5b). As expected, the posterior predictive check was somewhat better with the emulator, against which the posteriors were calibrated, than with MONARCH (*|z| <* 2 for 88.5% vs. 80.0% of pairs; coverage 88.1% vs. 78.5%).

**Fig. 5:**
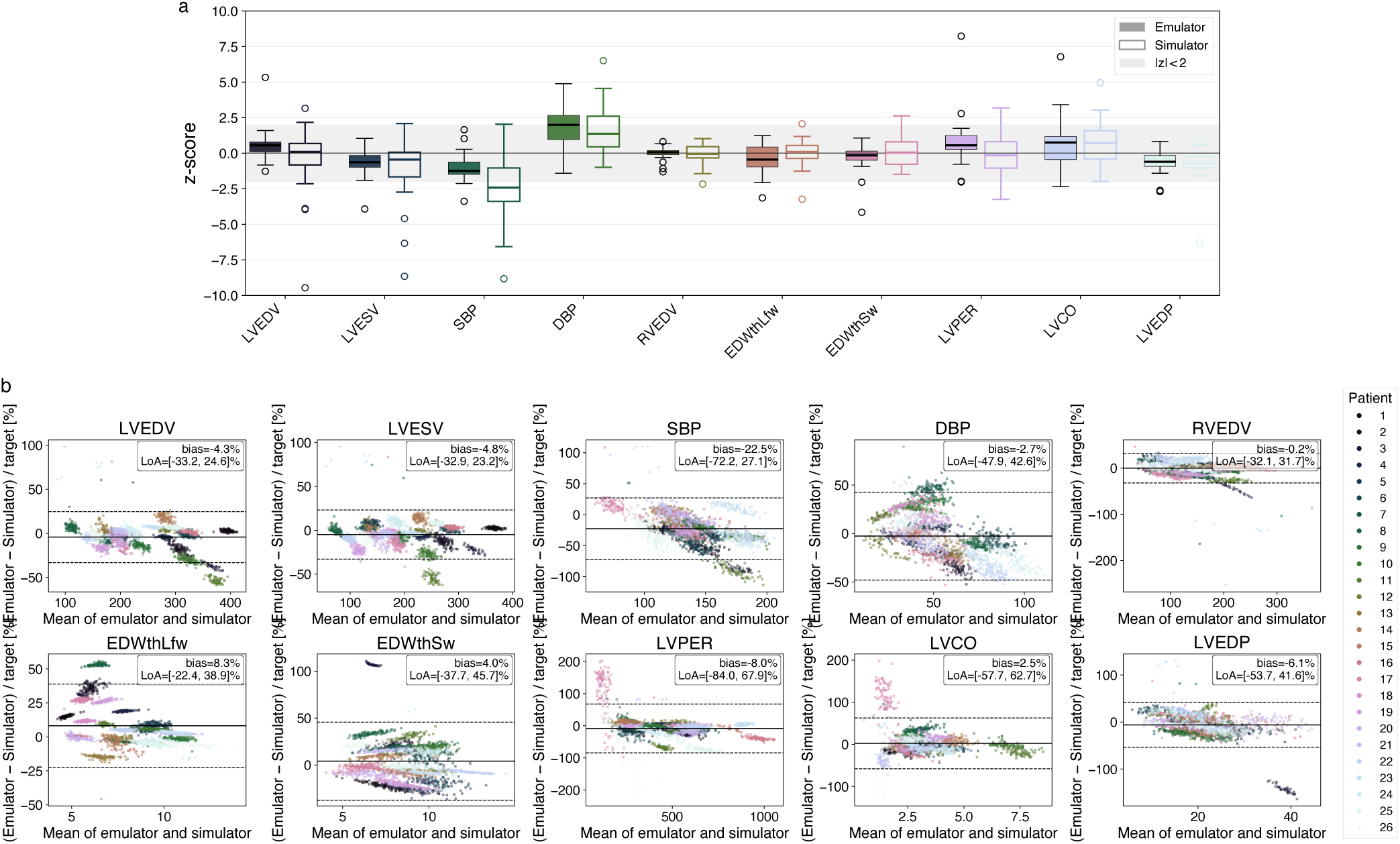
Emulator and simulator predictions at the calibrated posterior draws. **(a)** Standardized residuals *z* between each clinical target and the posterior predictive mean of the emulator (filled) and MONARCH (open) for the 26 patients; the grey band marks *|z| <* 2. **(b)** Bland–Altman plots of emulator versus MONARCH outputs at the posterior draws, as a percentage of the clinical target; solid lines, bias; dashed lines, 95% limits of agreement. Abbreviations as in Table 1.

The posterior draws produced patient-specific pressure–volume loops (Fig. 6), highlighting the diversity in the unseen patients calibrated with the population-level emulator. Posterior contraction and the correlations between parameters are reported in the Supplemental Material, Section S4.

**Fig. 6:**
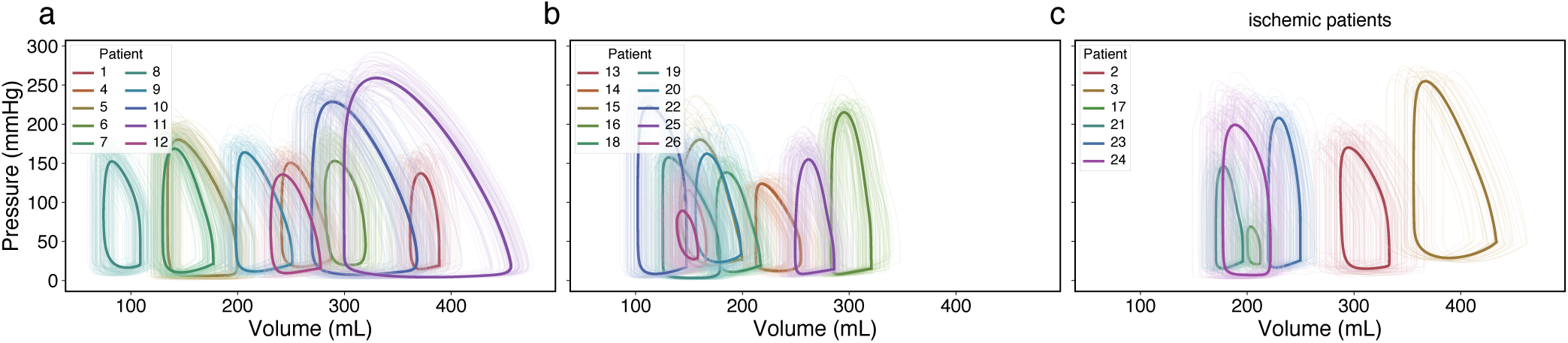
Calibrated digital twins reproduce patient-specific hemodynamics. Left-ventricular pressure–volume loops simulated with MONARCH at the posterior draws of each patient (thin lines) and their mean (thick lines), for the 20 patients without (a, b) and the six with (c) ischemic segments.

## 4 DISCUSSION

### 4.1 Summary of results

We developed a framework that calibrates a patient-specific cardiac model in about a minute, using a single emulator trained across a patient cohort. For patients the emulator had not seen, calibration took 59.7–63.0 s on a single CPU core, compared with an estimated 15 h with the simulator, and required no new simulations. The calibrated twins reproduced 80.0% of the clinical targets within two standard deviations, and most of the remaining misfit, concentrated in systolic blood pressure, reflects the reduced-order model rather than the emulator. Spending simulations on the real patients trained a more accurate emulator than adding virtual patients, and emulator accuracy was best for new patients whose clinical features were close to those in the training cohort.

### 4.2 Building on prior work of scalable surrogate models for cardiac digital twins

Surrogate modeling is now central to making large-scale cardiac and cardiovascular uncertainty quantification (UQ) tractable, with GP surrogates and multi-fidelity approaches delivering order-of-magnitude cost reductions in electrophysiology, mechanics, hemodynamics, and remodeling^14,16,47,58^. Earlier cardiac calibration work has shown that GP surrogates can both accelerate inverse problems and support global sensitivity analysis and inverse UQ for ventricular mechanics^19^ and atrial or whole-heart electrophysiology^59^. The central challenge we are addressing is that most emulator-based approaches for cardiac digital twins require the emulator to be trained for each individual patient.

Our population-emulator builds on two recent efforts to share the upfront cost of training simulations across a patient cohort. Rodero et al.^43^ reused simulations from previously calibrated patients to accelerate Bayesian history matching of new ones. Lanyon et al.^2^ proposed a latent-feature emulator, a single Gaussian process emulator for the whole cohort, and discrepancy emulators that learn how a new patient differs from the existing cohort. Their latent-feature emulator matched per-patient emulators at a fraction of the simulations for patients in the training cohort, but predicted left-out patients inconsistently, so that both of their methods still required additional simulations for every new patient. Both Rodero and Lanyon used three-dimensional models, using statistical shape models to represent geometry in a common latent space.

Our emulator follows Lanyon’s latent-feature design, but our latent features encode the clinical measurements that personalize the simulator (regional activation times, wall volumes and ischemia) rather than anatomy alone. This is facilitated by the use of the reduced-order TriSeg with MultiPatch heart model^38,40^ that explicitly includes these features in its formulation. Our generalization results agree with those of Lanyon: most held-out patients were predicted well (median NRMSE 11.0%), but error increased with distance from the cohort and a few isolated patients were predicted poorly (Section 3.2). What our results add is that this emulator accuracy suffices for Bayesian calibration that requires no additional simulations for a new patient, at the cost of lower accuracy than for patients included in training (emulator–MONARCH NRMSE 14.3% vs. 6.6% at the posterior draws). Patient-specific refinement, for example with a discrepancy emulator^2^, could potentially further improve this accuracy.

### 4.3 The choice of emulator design depends on the cohort and simulator

Although we demonstrated the framework on one cohort and one simulator, every new combination of the two requires the same design decisions we addressed in Section 3.1: how many simulations are needed, and whether they are better spent on the real patients or on synthetic patients placed between them. A different simulator and its free parameters, a larger or more heterogeneous cohort, or a different set of patient features may shift both the computational cost required and the balance between resampling and augmentation. The procedure we used to reach our configuration, however, generalizes, and follows the order of Section 3. First, tracing emulator accuracy against the computational cost for each training strategy sets the cost and the strategy. Second, leave-one-out cross-validation estimates accuracy for unseen patients and flags those far from the cohort. Third, checking the posteriors of held-out patients against the simulator exposes emulator error in the posterior region, where it matters for calibration.

For our cohort and reduced-order model of cardiac mechanics and hemodynamics, every output reached *R*^2^ *≥* 0.9 at *C* = 832 simulations, and spending a given computational cost on more simulations of the real patients (resampling) gave a more accurate emulator than adding synthetic patients (augmentation). This may seem counterintuitive for a cohort of only 26 patients (of whom only 6 had ischemic segments) which would seem set to benefit from enrichment, but could potentially be explained by how we generated synthetic patients.

There are various approaches to generate synthetic patients from an observed cohort: from interpolation between real patients, as here, to deep generative models such as variational autoencoders, which have generated virtual cardiac anatomies^50,60^ and virtual populations of digital twins^61^. Deep generative methods cover more of the feature space but need more data and make physiological plausibility harder to ensure; the models cited above were trained on 146 to more than 1,000 patients, far more than our 26. We therefore chose interpolation, which keeps every chimera within the convex hull of the cohort, but in doing so mostly adds support where the cohort is already dense rather than at its periphery, where errors were largest (Fig. 1). Sampling the latent space from a fitted Gaussian would extend beyond the hull at almost no cost, although the binary ischemia indicators that isolate the most peripheral patients are poorly described by a continuous distribution. Which augmentation strategy pays off will thus depend on the cohort and the simulator, but the procedure above can test it for any combination of the two.

### 4.4 Calibration misfit reflects the reduced-order model rather than the emulator

Most of the misfit that remained after calibration came from MONARCH rather than from the emulator. The calibrated twins overpredicted SBP (median *z* = *−*2.42) and tended to underpredict DBP (median *z* = +1.37), overestimating pulse pressure. Pulse pressure was already too wide on the emulator (Fig. 5a) and is set largely by stroke volume and systemic arterial capacitance, which we kept fixed at a literature value (*C*_as_; Appendix Table A2), so the sampler could not fit SBP and DBP together in many patients. Calibrating arterial capacitance would likely remove much of this misfit. The emulator added to the SBP misfit: at the posterior draws, the emulator–MONARCH difference in SBP averaged *−*22.5% of the target (95% limits of agreement *−*72.2% to +27.1%), so the sampler accepted parameters whose simulated SBP was higher than the emulator predicted. For DBP, by contrast, the bias was only *−*2.7%, so its misfit reflects the model rather than the emulator. Finally, 13.1% of the posterior draws did not produce a valid MONARCH simulation, most of them in 7 patients, and a few produced degenerate pressure– volume loops, indicating posterior regions where the reduced-order model is not physiologically valid. A different simulator may fit these patients better or worse; the framework itself does not depend on the simulator, and the posterior predictive check against it allows for distinguishing model error from emulator error.

### 4.5 Choice of sampler and alternative Bayesian methods

The choice of MCMC sampler matters: samplers differ in how efficiently they explore a posterior and in how reliably they capture correlated or multimodal posteriors. A cheap emulator makes this choice affordable rather than fixing it, so the sampler can be chosen for the model and data at hand. We used the gradient-free DEMetropolisZ sampler, which needs no tuning of step sizes or mass matrices and parallelizes over chains. Posteriors converged in about a minute for every patient; five patients fell slightly short of the strictest convergence thresholds, which longer chains could resolve at little cost. Because the GP posterior mean is differentiable, gradient-based samplers can be used as well: within cardiac modeling, Hamiltonian Monte Carlo (HMC) has been shown to recover parameter distributions efficiently for action potential models and compact cardiovascular surrogates^62–64^, and physics-informed GP calibration frameworks rely on it for differential-equation models, including cardiovascular ODEs^65^.

Alternative Bayesian strategies trade accuracy for cost: Bayesian history matching rules out implausible parameter regions rather than estimating a posterior^12,43^, ensemble Kalman filters approximate the posterior as Gaussian^59^, and approximate Bayesian computation with sequential Monte Carlo avoids an explicit likelihood^62^. With a cheap and pre-trained emulator, full MCMC is affordable, and the shape of the resulting posteriors is itself informative: the targets constrained all seven parameters in most patients, stressed blood volume least, and the strongest posterior correlation, a trade-off between the left free wall and septal reference areas (mean *r* = *−*0.54), is consistent with both walls jointly setting left-ventricular cavity size (Supplemental Material, Section S4).

### 4.6 Strengths and clinical implications

The principal strength of the proposed framework is the combination of *rigor* and *speed*. Calibration for patients the emulator was not trained on is fast enough for clinical workflows, and it remains fully Bayesian: measurement uncertainty enters the likelihood explicitly, and posterior predictive checks against the simulator quantify the remaining emulator error. Driving the same sampler directly with MONARCH would have taken an estimated 15 h per patient (120,000 model evaluations, i.e. 4 chains of 20,000 draws plus 10,000 tuning steps, at 0.44 s each), and far longer for high-fidelity models whose simulations take minutes to hours each. Admitting a new patient requires only computing their latent representation from the patient features: the trained emulator then predicts their outputs without retraining, which is the central advantage of a cohort-wide emulator over per-patient surrogates.

For patients referred for CRT, for example, like those in our cohort, this opens several previously infeasible workflows. Pre-procedural planning could include patient-specific simulations of candidate pacing strategies, such as conventional biventricular pacing versus conduction system pacing, as well as lead positions or pacing protocols using the calibrated baseline twin as the starting point^7,29,66^. The framework also provides a natural foundation for treatment-response prediction: a calibrated baseline twin can be coupled with a forward model of post-CRT remodeling to provide probabilistic predictions of response^25^, with credible intervals that propagate from the baseline calibration^14^. This is complementary to the growing body of work that predicts CRT response with machine learning from clinical variables, imaging, and ECG^67–70^: rather than predicting response directly, the framework infers interpretable parameter distributions with uncertainty for each patient, which could serve as mechanistic features for such models.

Beyond CRT, our emulator framework can be used wherever patient-specific models must be calibrated to scalar biomarkers in small cohorts, a setting common in but not limited to precision cardio-vascular medicine^1,3^. It could further extend from our reduced-order model to high-fidelity models by encoding patient-specific geometry by the modes of a statistical shape model^2,43^.

### 4.7 Limitations

Several limitations should be considered when interpreting these results.

First, the test cohort of 26 patients is small, exacerbated by only 6 ischemic patients, and the emulator’s accuracy for a new patient depends on how well the cohort represents that patient. This also makes this cohort a suitable stress test: isolated patients, the ischemic ones in particular, had few close neighbors to learn from, a challenge that would not have been posed by larger or more homogeneous cohorts. The most peripheral patients remained the hardest to predict, and chimera augmentation, which cannot leave the convex hull of the observed features, did not weaken the dependence of perpatient error on distance from the cohort (Fig. 3, Section 4.3). Prior work on cohort emulation likewise recommends refined or patient-specific emulators for patients far from the training region^2,43^. A new patient’s latent distance to the cohort can flag likely extrapolation before calibration, but it does not remedy it.

Second, several design choices were set from cohort-level diagnostics rather than optimized jointly: the Morris screening threshold, the number of principal components, the chimera count and the computational cost. Other choices may suit other models or cohorts, and adaptive, multi-wave refinement of the training design or multi-fidelity strategies may reach the same accuracy at lower upfront cost^16,58^.

Third, the MONARCH simulator is a reduced-order representation of the heart and circulation that omits three-dimensional geometry, regional fiber architecture and detailed valve mechanics. The framework is agnostic to the forward model, but each calibrated twin is only as faithful as that model, and our focus here is on the quality of the calibration rather than limitations of the MONARCH. The systematic overestimation of pulse pressure (Section 4.4) is one sign of such model-form error; parameters fixed at literature values, such as systemic arterial compliance, may need to be calibrated in future applications.

Fourth, the posterior predictive intervals were somewhat narrower than nominal: the 95% interval, including measurement noise, contained the clinical target for 78.5% of patient–target pairs, with most misses in SBP. The likelihood treats the model, the emulator and the patient features as exact. It includes no model-discrepancy term and not the emulator’s predictive variance, so posteriors may be too narrow or shifted where the model or emulator is inaccurate, as both were for SBP; adding the emulator variance to the likelihood is a direct remedy.

Fifth, the patient features were prescribed as exact, although each carries measurement error. Of these, only the activation times required DENSE MRI, which is not part of routine clinical imaging; they could instead be estimated from the ECG, for example by scaling an electrophysiology model of the activation sequence to the QRS duration.

Sixth, two of the calibration targets are imperfect proxies. LVEDP was estimated from NYHA class rather than measured (Appendix A2), because invasive left-ventricular catheterization is not routinely performed in patients referred for CRT, and LVCO is defined differently in the data (volumetric) and in the model (forward flow through the aortic valve), two definitions that differed by more than 10% for about a quarter of the posterior draws (Section 2.1); calibrating against MONARCH’s volumetric LVCO would remove this mismatch. Such incomplete and indirect measurements are typical of routine clinical data, however, and the framework accommodates them: each target enters the likelihood with its own uncertainty, so that less certain targets such as LVEDP weigh less in the calibration.

### 4.8 Future perspectives

We set out to calibrate patient-specific cardiac digital twins within minutes of new data becoming available, with an honest estimate of their uncertainty. The framework presented here does so by training a single emulator across a patient cohort, which exploits the structure the patients share: each held-out patient was calibrated with an emulator trained on the other 25 patients in about a minute on a single CPU core, without any new simulations, and the calibrated twins reproduced 80.0% of the clinical targets within two standard deviations. Such twins could serve as the starting point for patient-specific predictions, such as the response to different CRT strategies.

More broadly, training an emulator once across a cohort and calibrating each new patient without new simulations offers a general recipe for accelerating mechanistic patient-specific modeling. The simulator, the surrogate and the sampler are interchangeable, and the latent encoding accepts whatever patient features personalize the simulator, from the regional measurements used here to the shape modes that describe patient-specific geometry in high-fidelity models^2,43^. The best training design will differ for each combination of cohort, surrogate model, and simulator, but can be determined by following the procedure we followed here.

By removing the need for new simulations for every new patient, the framework brings uncertainty-aware cardiac digital twins a step closer to routine clinical workflows.

## 5 DATA AND SOFTWARE AVAILABILITY

These methodologies and results will be made publicly available on GitHub upon publication of this paper at https://github.com/BeatLabUCI/. Curated Jupyter notebooks with extensive documentation are included to reproduce all results in this manuscript, as well as tutorial notebooks to help others use our code to calibrate their own models. Further inquiries can be addressed to the corresponding author.

The clinical data supporting these findings were collected under protocols approved by the University of Virginia Institutional Review Board and are not publicly available because patient consent did not cover open redistribution. De-identified derived data (calibration targets and patient features) are available from the corresponding author on reasonable request, subject to a data use agreement.

## ACKNOWLEDGMENTS

This research was funded by the National Heart, Lung, and Blood Institute of the National Institutes of Health (R01HL159945).

## AUTHORS CONTRIBUTIONS

C.E.J.: Methodology, Software, Investigation, Formal analysis, Writing – original draft, Writing – review & editing.

F.P.: Methodology, Software, Investigation, Formal analysis, Writing – review & editing.

D.J.B.: Data curation, Methodology, Investigation, Writing – review & editing.

J.H.B.: Methodology, Investigation, Writing – review & editing.

Z.H.: Methodology, Software, Investigation.

I.A.: Methodology, Investigation. L.L.: Methodology, Investigation.

K.C.B.: Data curation, Methodology, Investigation, Writing – review & editing, Funding acquisition, Supervision.

P.J.A.O.: Conceptualization, Methodology, Software, Investigation, Formal analysis, Writing – review & editing, Funding acquisition, Supervision.

## DECLARATIONS

C.E.J. is an employee of Edwards Lifesciences. The research presented here is independent of her employment and was conducted as part of her PhD work at UC Irvine. All other authors declare no conflicts of interest.

## CITATION DIVERSITY STATEMENT

Papers by women and other underrepresented scholars are undercited relative to their share of the field^71,72^. We therefore obtained the predicted gender of the first and last author of each reference from databases that store the probability of a first name being carried by a woman^73,74^. Excluding self-citations to the first and last authors of this paper and the references in this statement, our references contain 2% woman(first)/woman(last), 20% man/woman, 18% woman/man and 60% man/man. We also obtained the predicted racial/ethnic category of the first and last author of each reference from their first and last names^75^. By this measure, our references contain 9% author of color(first)/author of color(last), 14% white author/author of color, 26% author of color/white author and 51% white author/white author. These methods are limited in that names may not indicate gender or racial/ethnic identity, and they cannot account for intersex, non-binary or transgender people, or for Indigenous and mixed-race authors.

## A1 MODEL PARAMETERS

**Table A1:**
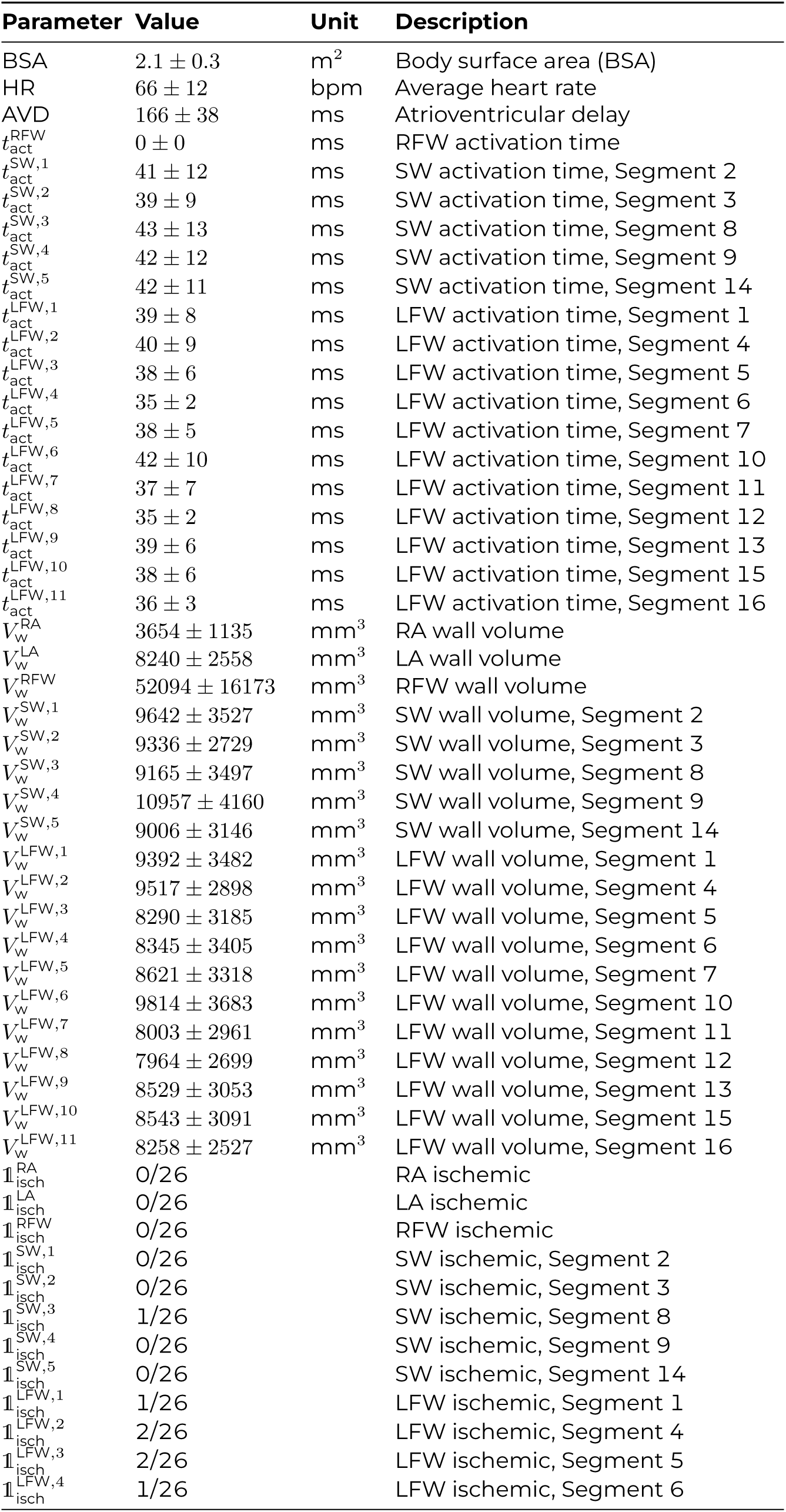
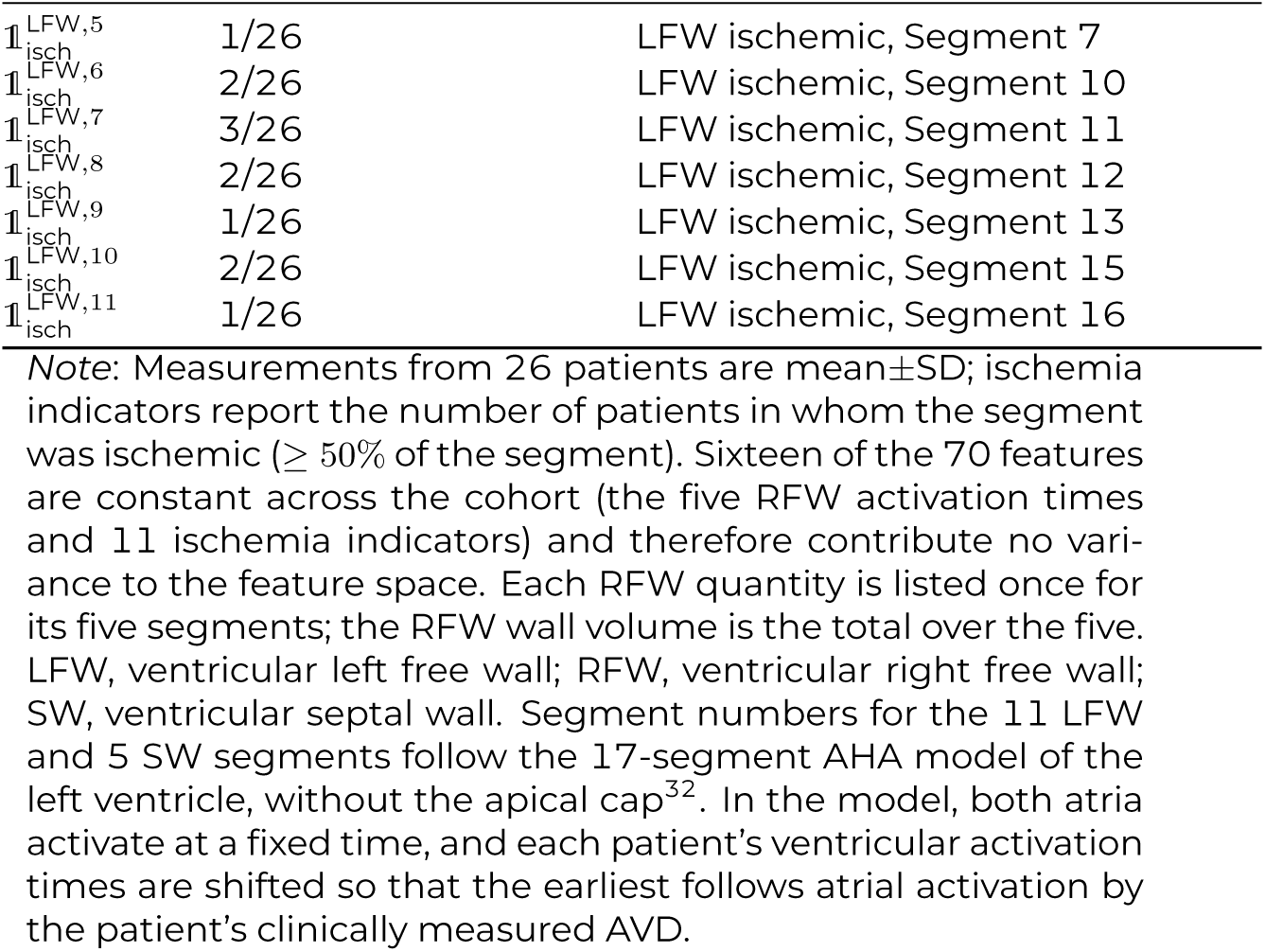
Clinical measurements in set *D* of the 26 patients, prescribed directly as model inputs (Eq. 1). All entries except HR enter the patient feature vector *F* used by the emulator.

| Parameter | Value | Unit | Description |
|---|---|---|---|
| BSA | 2.1 ± 0.3 | m<sup>2</sup> | Body surface area (BSA) |
| HR | 66 ± 12 | bpm | Average heart rate |
| AVD | 166 ± 38 | ms | Atrioventricular delay |
| *t*<sub>act</sub><sup>RFW</sup> | 0 ± 0 | ms | RFW activation time |
| *t*<sub>act</sub><sup>SW,1</sup> | 41 ± 12 | ms | SW activation time, Segment 2 |
| *t*<sub>act</sub><sup>SW,2</sup> | 39 ± 9 | ms | SW activation time, Segment 3 |
| *t*<sub>act</sub><sup>SW,3</sup> | 43 ± 13 | ms | SW activation time, Segment 8 |
| *t*<sub>act</sub><sup>SW,4</sup> | 42 ± 12 | ms | SW activation time, Segment 9 |
| *t*<sub>act</sub><sup>SW,5</sup> | 42 ± 11 | ms | SW activation time, Segment 14 |
| *t*<sub>act</sub><sup>LFW,1</sup> | 39 ± 8 | ms | LFW activation time, Segment 1 |
| *t*<sub>act</sub><sup>LFW,2</sup> | 40 ± 9 | ms | LFW activation time, Segment 4 |
| *t*<sub>act</sub><sup>LFW,3</sup> | 38 ± 6 | ms | LFW activation time, Segment 5 |
| *t*<sub>act</sub><sup>LFW,4</sup> | 35 ± 2 | ms | LFW activation time, Segment 6 |
| *t*<sub>act</sub><sup>LFW,5</sup> | 38 ± 5 | ms | LFW activation time, Segment 7 |
| *t*<sub>act</sub><sup>LFW,6</sup> | 42 ± 10 | ms | LFW activation time, Segment 10 |
| *t*<sub>act</sub><sup>LFW,7</sup> | 37 ± 7 | ms | LFW activation time, Segment 11 |
| *t*<sub>act</sub><sup>LFW,8</sup> | 35 ± 2 | ms | LFW activation time, Segment 12 |
| *t*<sub>act</sub><sup>LFW,9</sup> | 39 ± 6 | ms | LFW activation time, Segment 13 |
| *t*<sub>act</sub><sup>LFW,10</sup> | 38 ± 6 | ms | LFW activation time, Segment 15 |
| *t*<sub>act</sub><sup>LFW,11</sup> | 36 ± 3 | ms | LFW activation time, Segment 16 |
| *V*<sub>w</sub><sup>RA</sup> | 3654 ± 1135 | mm<sup>3</sup> | RA wall volume |
| *V*<sub>w</sub><sup>LA</sup> | 8240 ± 2558 | mm<sup>3</sup> | LA wall volume |
| *V*<sub>w</sub><sup>RFW</sup> | 52094 ± 16173 | mm<sup>3</sup> | RFW wall volume |
| *V*<sub>w</sub><sup>SW,1</sup> | 9642 ± 3527 | mm<sup>3</sup> | SW wall volume, Segment 2 |
| *V*<sub>w</sub><sup>SW,2</sup> | 9336 ± 2729 | mm<sup>3</sup> | SW wall volume, Segment 3 |
| *V*<sub>w</sub><sup>SW,3</sup> | 9165 ± 3497 | mm<sup>3</sup> | SW wall volume, Segment 8 |
| *V*<sub>w</sub><sup>SW,4</sup> | 10957 ± 4160 | mm<sup>3</sup> | SW wall volume, Segment 9 |
| *V*<sub>w</sub><sup>SW,5</sup> | 9006 ± 3146 | mm<sup>3</sup> | SW wall volume, Segment 14 |
| *V*<sub>w</sub><sup>LFW,1</sup> | 9392 ± 3482 | mm<sup>3</sup> | LFW wall volume, Segment 1 |
| *V*<sub>w</sub><sup>LFW,2</sup> | 9517 ± 2898 | mm<sup>3</sup> | LFW wall volume, Segment 4 |
| *V*<sub>w</sub><sup>LFW,3</sup> | 8290 ± 3185 | mm<sup>3</sup> | LFW wall volume, Segment 5 |
| *V*<sub>w</sub><sup>LFW,4</sup> | 8345 ± 3405 | mm<sup>3</sup> | LFW wall volume, Segment 6 |
| *V*<sub>w</sub><sup>LFW,5</sup> | 8621 ± 3318 | mm<sup>3</sup> | LFW wall volume, Segment 7 |
| *V*<sub>w</sub><sup>LFW,6</sup> | 9814 ± 3683 | mm<sup>3</sup> | LFW wall volume, Segment 10 |
| *V*<sub>w</sub><sup>LFW,7</sup> | 8003 ± 2961 | mm<sup>3</sup> | LFW wall volume, Segment 11 |
| *V*<sub>w</sub><sup>LFW,8</sup> | 7964 ± 2699 | mm<sup>3</sup> | LFW wall volume, Segment 12 |
| *V*<sub>w</sub><sup>LFW,9</sup> | 8529 ± 3053 | mm<sup>3</sup> | LFW wall volume, Segment 13 |
| *V*<sub>w</sub><sup>LFW,10</sup> | 8543 ± 3091 | mm<sup>3</sup> | LFW wall volume, Segment 15 |
| *V*<sub>w</sub><sup>LFW,11</sup> | 8258 ± 2527 | mm<sup>3</sup> | LFW wall volume, Segment 16 |
| 𝟙<sub>isch</sub><sup>RA</sup> | 0/26 |  | RA ischemic |
| 𝟙<sub>isch</sub><sup>LA</sup> | 0/26 |  | LA ischemic |
| 𝟙<sub>isch</sub><sup>RFW</sup> | 0/26 |  | RFW ischemic |
| 𝟙<sub>isch</sub><sup>SW,1</sup> | 0/26 |  | SW ischemic, Segment 2 |
| 𝟙<sub>isch</sub><sup>SW,2</sup> | 0/26 |  | SW ischemic, Segment 3 |
| 𝟙<sub>isch</sub><sup>SW,3</sup> | 1/26 |  | SW ischemic, Segment 8 |
| 𝟙<sub>isch</sub><sup>SW,4</sup> | 0/26 |  | SW ischemic, Segment 9 |
| 𝟙<sub>isch</sub><sup>SW,5</sup> | 0/26 |  | SW ischemic, Segment 14 |
| 𝟙<sub>isch</sub><sup>LFW,1</sup> | 1/26 |  | LFW ischemic, Segment 1 |
| 𝟙<sub>isch</sub><sup>LFW,2</sup> | 2/26 |  | LFW ischemic, Segment 4 |
| 𝟙<sub>isch</sub><sup>LFW,3</sup> | 2/26 |  | LFW ischemic, Segment 5 |
| 𝟙<sub>isch</sub><sup>LFW,4</sup> | 1/26 |  | LFW ischemic, Segment 6 |
| 𝟙<sub>isch</sub><sup>LFW,5</sup> | 1/26 |  | LFW ischemic, Segment 7 |
| 𝟙<sub>isch</sub><sup>LFW,6</sup> | 2/26 |  | LFW ischemic, Segment 10 |
| 𝟙<sub>isch</sub><sup>LFW,7</sup> | 3/26 |  | LFW ischemic, Segment 11 |
| 𝟙<sub>isch</sub><sup>LFW,8</sup> | 2/26 |  | LFW ischemic, Segment 12 |
| 𝟙<sub>isch</sub><sup>LFW,9</sup> | 1/26 |  | LFW ischemic, Segment 13 |
| 𝟙<sub>isch</sub><sup>LFW,10</sup> | 2/26 |  | LFW ischemic, Segment 15 |
| 𝟙<sub>isch</sub><sup>LFW,11</sup> | 1/26 |  | LFW ischemic, Segment 16 |

*Note*: Measurements from 26 patients are mean±SD; ischemia indicators report the number of patients in whom the segment was ischemic (≥ 50% of the segment). Sixteen of the 70 features are constant across the cohort (the five RFW activation times and 11 ischemia indicators) and therefore contribute no vari-ance to the feature space. Each RFW quantity is listed once for its five segments; the RFW wall volume is the total over the five. LFW, ventricular left free wall; RFW, ventricular right free wall; SW, ventricular septal wall. Segment numbers for the 11 LFW and 5 SW segments follow the 17-segment AHA model of the left ventricle, without the apical cap<sup>32</sup>. In the model, both atria activate at a fixed time, and each patient's ventricular activation times are shifted so that the earliest follows atrial activation by the patient's clinically measured AVD.

**Table A2:** Constant model parameters in set *C* that are set equal for the whole patient population.

| Model | Parameter | Value | Unit | Source | Description |
|---|---|---|---|---|---|
| Constitutive (active) | *L*<sub>s,ref</sub> | 2.0 | *µ*m | <sup>38</sup> | Reference sarcomere length |
|  | *L*<sub>se,iso</sub> | 0.04 | *µ*m | <sup>38</sup> | Series elastic element length |
|  | *L*<sub>sc,0</sub> | 1.51 | *µ*m | <sup>38</sup> | Contractile element length at zero stress |
|  | *v*<sub>max,v</sub> | 7 | *µ*m · *s*<sup>−1</sup> | <sup>38</sup> | Maximum ventricular sarcomere shortening velocity |
|  | *v*<sub>max,a</sub> | 14 | *µ*m · *s*<sup>−1</sup> | <sup>14</sup> | Maximum atrial sarcomere shortening velocity |
|  | *s*<sub>fact,a</sub> | 0.06 | MPa | <sup>14</sup> | Atrial active stress coefficient |
|  | *τ*<sub>R,v</sub> | 0.25·*t*<sub>ad</sub> | ms | <sup>14</sup> | Factor scaling ventricular contraction rise time |
|  | *τ*<sub>D,v</sub> | 0.25·*t*<sub>ad</sub> | ms | <sup>14</sup> | Factor scaling ventricular contraction decay time |
|  | *τ*<sub>R,a</sub> | 0.40·*t*<sub>ad,a</sub> | ms | <sup>14</sup> | Factor scaling atrial contraction rise time |
|  | *τ*<sub>D,a</sub> | 0.40·*t*<sub>ad,a</sub> | ms | <sup>14</sup> | Factor scaling atrial contraction decay time |
|  | *t*<sub>ad,a</sub> | 80 | ms | <sup>14</sup> | Atrial contraction duration |
| Constitutive (passive) | *c*<sub>1,v</sub> | 1·10<sup>−5</sup> | MPa | <sup>14</sup> | Ventricular passive stress coefficient |
|  | *c*<sub>3,v</sub> | 5.23·10<sup>−4</sup> | MPa | <sup>14</sup> | Ventricular passive stress coefficient |
|  | *c*<sub>4,v</sub> | 10 |  | <sup>14</sup> | Ventricular passive stress coefficient |
|  | *c*<sub>1,a</sub> | 1.60·10<sup>−3</sup> | MPa | <sup>14</sup> | Atrial passive stress coefficient |
|  | *c*<sub>3,a</sub> | 1.7·10<sup>−5</sup> | MPa | <sup>14</sup> | Atrial passive stress coefficient |
|  | *c*<sub>4,a</sub> | 10 |  | <sup>14</sup> | Atrial passive stress coefficient |
| Circulation | *R*<sub>cs</sub> | 0.090 | mmHg·s·mL<sup>−1</sup> | <sup>14</sup> | Systemic characteristic resistance |
|  | *R*<sub>vs</sub> | 0.015 | mmHg·s·mL<sup>−1</sup> | <sup>24</sup> | Systemic venous resistance |
|  | *R*<sub>cp</sub> | 0.020 | mmHg·s·mL<sup>−1</sup> | <sup>24</sup> | Pulmonary characteristic resistance |
|  | *R*<sub>ap</sub> | 0.300 | mmHg·s·mL<sup>−1</sup> | <sup>14</sup> | Pulmonary arterial resistance |
|  | *R*<sub>vp</sub> | 0.015 | mmHg·s·mL<sup>−1</sup> | <sup>24</sup> | Pulmonary venous resistance |
|  | *R*<sub>av</sub> | 0.025 | mmHg·s·mL<sup>−1</sup> | <sup>14</sup> | Atrioventricular resistance |
|  | *C*<sub>as</sub> | 0.36 | mL·mmHg<sup>−1</sup> | <sup>24</sup> | Systemic arterial capacitance |
|  | *C*<sub>vs</sub> | 70 | mL·mmHg<sup>−1</sup> | <sup>24</sup> | Systemic venous capacitance |
|  | *C*<sub>ap</sub> | 13 | mL·mmHg<sup>−1</sup> | <sup>24</sup> | Pulmonary arterial capacitance |
|  | *C*<sub>vp</sub> | 8 | mL·mmHg<sup>−1</sup> | <sup>24</sup> | Pulmonary venous capacitance |
| Cardiac geometry | *A*<sub>m,ref,RA</sub> | 4840 | mm<sup>2</sup> | <sup>40</sup> | RA midwall reference area |
|  | *A*<sub>m,ref,LA</sub> | 6679 | mm<sup>2</sup> | <sup>40</sup> | LA midwall reference area |

## A2 ESTIMATION OF LVEDP

Left ventricular end-diastolic pressure was not measured invasively in this cohort. We approximated it by pulmonary capillary wedge pressure (PCWP), which approximates LVEDP in the absence of mitral stenosis, and assigned each patient the PCWP associated with their NYHA functional class. To estimate these values, we reviewed clinical studies of patients with heart failure that grouped participants by NYHA class and reported PCWP measured by catheterization, and collected data from six studies^76–81^. For each NYHA class we calculated the sample-size-weighted mean and standard deviation of PCWP across studies, assuming zero covariance between studies, and rounded up to the next integer (Fig. A1). This gave a mean PCWP of 15 mmHg for NYHA class II (109 individuals in total) and 23 mmHg for class III (71 individuals), with weighted standard deviations of 4.67 and 4.62 mmHg, which we rounded up to 5 mmHg for both classes. PCWP increased with NYHA class (*R*^2^ = 0.64). In our cohort, 20 patients were in class II and 6 in class III, so LVEDP took only two values (Table 1), and the cohort median coincides with the minimum. We used the 5 mmHg standard deviation as the measurement uncertainty of LVEDP; because it reflects the spread of PCWP within a NYHA class rather than a patient-specific measurement, LVEDP is the least certain of the ten targets.

**Fig. A1:**
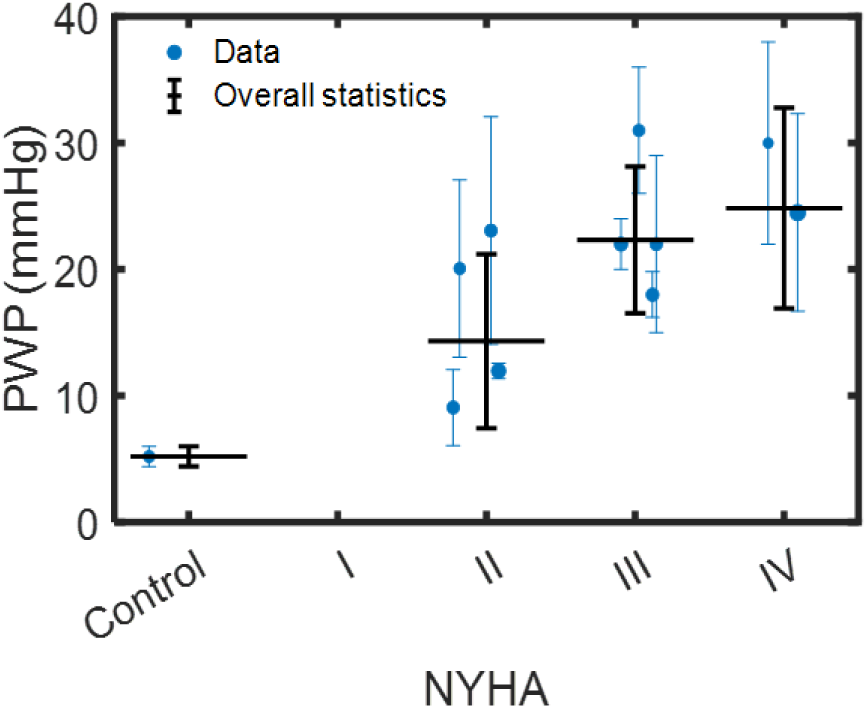
Pulmonary capillary wedge pressure increases with NYHA functional class. PCWP (PWP) by NYHA class reported in six clinical studies of patients with heart failure^76–81^. Blue points and error bars show the values reported by each study; black bars show the sample-size-weighted mean and standard deviation across studies (overall statistics), from which the LVEDP targets of 15 mmHg (class II) and 23 mmHg (class III) were derived.

## REFERENCES

[1] P. Thangaraj, S. Benson, E. Oikonomou, F. Asselbergs, and R. Khera. “Cardiovascular care with digital twin technology in the era of generative artificial intelligence”. In: European Heart Journal (2024). doi: 10.1093/eurheartj/ehae619.

[2] C. W. Lanyon, C. Rodero, A. Qayyum, T. M. Baptiste, S. A. Niederer, and R. D. Wilkinson. “Weaving the Digital Tapestry: Methods for Emulating Cohorts of Cardiac Digital Twins Using Gaussian Processes”. In: sAnnals of Biomedical Engineering 54.2 (Feb. 2026), pp. 514–530. doi: 10.1007/s10439-025-03890-0.

[3] M. Colebank, P. Oomen, C. Witzenburg, A. Grosberg, D. Beard, D. Husmeier, M. Olufsen, and N. Chesler. “Guidelines for mechanistic modeling and analysis in cardiovascular research”. In: American Journal of Physiology - Heart and Circulatory Physiology 327 (2024), H473–H503. doi: 10.1152/ajpheart.00766.2023.

[4] E. Zappon et al. “An efficient end-to-end computational framework for the generation of ECG calibrated volumetric models of human atrial electrophysiology”. In: Medical Image Analysis 107 (2025), p. 103822. doi: 10.1016/j.media.2025.103822.

[5] A. Jung, M. Gsell, C. Augustin, and G. Plank. “An Integrated Workflow for Building Digital Twins of Cardiac Electromechanics — A Multi-Fidelity Approach for Personalising Active Mechanics”. In: Mathematics 10 (2022). doi: 10.3390/math10050823.

[6] J. Xu and F. Wang. “Cardiac Mechano-Electrical-Fluid Interaction: A Brief Review of Recent Advances”. In: Eng (2025). doi: 10.3390/eng6080168.

[7] T. Koopsen et al. “Virtual pacing of a patient’s digital twin to predict left ventricular reverse remodelling after cardiac resynchronization therapy”. In: Europace 26.1 (2024), euae009. doi: 10.1093/europace/euae009.

[8] J. Chrispin, A. Prakosa, E. Kholmovski, A. Kolandaivelu, A. Barcelon, K. N. Aronis, R. D. Berger, H. Calkins, and N. Trayanova. “Digital Twin–Guided Ablation for Ventricular Tachycardia”. en. In: New England Journal of Medicine 394.13 (Apr. 2026), pp. 1345–1347. doi: 10.1056/NEJMc2517822.

[9] N. Trayanova, A. Lyon, J. Shade, and J. Heijman. “Computational modeling of cardiac electrophysiology and arrhythmogenesis: toward clinical translation”. In: Physiological Reviews 104 (2023), pp. 1265–1333. doi: 10.1152/physrev.00017.2023.

[10] O. Jaffery, L. Melki, G. Slabaugh, W. Good, and C. Roney. “A Review of Personalised Cardiac Computational Modelling Using Electroanatomical Mapping Data”. In: Arrhythmia & Electrophysiology Review 13 (2024). doi: 10.15420/aer.2023.25.

[11] W. H. Chen, G. R. Gavalas, J. H. Seinfeld, and M. L. Wasserman. “A New Algorithm for Automatic History Matching”. In: Society of Petroleum Engineers Journal 14.06 (1974), pp. 593–608. doi: 10.2118/4545-pa.

[12] I. Vernon, M. Goldstein, and R. Bower. “Galaxy Formation: Bayesian History Matching for the Observable Universe”. In: Statistical Science 29.1 (2014). doi: 10.1214/12-sts412. eprint: 1405.4976.

[13] S. Longobardi, A. Lewalle, S. Coveney, I. Sjaastad, E. K. S. Espe, W. E. Louch, C. J. Musante, A. Sher, and S. A. Niederer. “Predicting left ventricular contractile function via Gaussian process emulation in aortic-banded rats”. In: Philosophical Transactions of the Royal Society A 378.2173 (2020), p. 20190334. doi: 10.6084/m9.figshare.12300875.v2.

[14] C. E. Jones and P. J. Oomen. “Synergistic biophysics and machine learning modeling to rapidly predict cardiac growth probability”. en. In: Computers in Biology and Medicine 184 (Jan. 2025), p. 109323. doi: 10.1016/j.compbiomed.2024.109323.

[15] C. Corrado et al. “Quantifying the impact of shape uncertainty on predicted arrhythmias”. In: Computers in Biology and Medicine 153 (2023), p. 106528. doi: 10.1016/j.compbiomed.2022.106528.

[16] C. Fleeter, G. Geraci, D. Schiavazzi, A. Kahn, and A. Marsden. “Multilevel and multifidelity uncertainty quantification for cardiovascular hemodynamics”. In: Computer Methods in Applied Mechanics and Engineering 365 (2019). doi: 10.1016/j.cma.2020.113030.

[17] G. Mirams, P. Pathmanathan, R. Gray, P. Challenor, and R. Clayton. “Uncertainty and variability in computational and mathematical models of cardiac physiology”. In: The Journal of Physiology 594 (2016), pp. 6833–6847. doi: 10.1113/jp271671.

[18] S. Pagani and A. Manzoni. “Enabling forward uncertainty quantification and sensitivity analysis in cardiac electrophysiology by reduced order modeling and machine learning”. In: International Journal for Numerical Methods in Biomedical Engineering 37 (2021). doi: 10.1002/cnm.3450.

[19] A. Lazarus, D. Dalton, D. Husmeier, and H. Gao. “Sensitivity analysis and inverse uncertainty quantification for the left ventricular passive mechanics”. In: Biomechanics and Modeling in Mechanobiology 21 (2022), pp. 953–982. doi: 10.1007/s10237-022-01571-8.

[20] C. Lei et al. “Considering discrepancy when calibrating a mechanistic electrophysiology model”. In: Philosophical Transactions of the Royal Society A: Mathematical, Physical and Engineering Sciences 378 (2020). doi: 10.1098/rsta.2019.0349.

[21] A. Marquis, A. Arnold, C. Dean-Bernhoft, B. Carlson, and M. Olufsen. “Practical identifiability and uncertainty quantification of a pulsatile cardiovascular model”. In: Mathematical Biosciences 304 (2017), pp. 9–24. doi: 10.1016/j.mbs.2018.07.001.

[22] B. Winkler, C. Nagel, N. Farchmin, S. Heidenreich, A. Loewe, O. Dössel, and M. Bär. “Global Sensitivity Analysis and Uncertainty Quantification for Simulated Atrial Electrocardiograms”. In: Metrology (2022). doi: 10.3390/metrology3010001.

[23] M. Salvador, F. Kong, M. Peirlinck, D. W. Parker, H. Chubb, A. M. Dubin, and A. L. Marsden. Digital twinning of cardiac electrophysiology for congenital heart disease. en. preprint. Bioengineering, Nov. 2023. doi: 10.1101/2023.11.27.568942.

[24] D. Burkhoff and J. V. Tyberg. “Why does pulmonary venous pressure rise after onset of LV dysfunction: a theoretical analysis”. In: American Journal of Physiology-Heart and Circulatory Physiology 265.5 (1993), H1819–H1828. doi: 10.1152/ajpheart.1993.265.5.h1819.

[25] P. J. A. Oomen, T.-K. N. Phung, S. H. Weinberg, K. C. Bilchick, and J. W. Holmes. “A rapid electromechanical model to predict reverse remodeling following cardiac resynchronization therapy”. In: Biomechanics and Modeling in Mechanobiology 21.1 (2022), pp. 231–247. doi: 10.1007/s10237-021-01532-7.

[26] W. P. Santamore and D. Burkhoff. “Hemodynamic consequences of ventricular interaction as assessed by model analysis”. In: American Journal of Physiology-Heart and Circulatory Physiology 260.1 (1991), H146–H157. doi: 10.1152/ajpheart.1991.260.1.h146.

[27] J. Walmsley, P. R. Huntjens, F. W. Prinzen, T. Delhaas, and J. Lumens. “Septal flash and septal rebound stretch have different underlying mechanisms”. In: American Journal of Physiology-Heart and Circulatory Physiology 310.3 (2016), H394–H403. doi: 10.1152/ajpheart.00639.2015.

[28] W. L. Maughan, K. Sunagawa, D. Burkhoff, W. L. G. Jr, W. C. Hunter, and K. Sagawa. “Effect of heart rate on the canine end-systolic pressure-volume relationship.” In: Circulation 72.3 (1985), pp. 654–659. doi: 10.1161/01.cir.72.3.654.

[29] M. Taconné, V. Rolle, E. Galli, K. Owashi, A. Wazzan, E. Donal, and A. Hernández. “Characterization of cardiac resynchronization therapy response through machine learning and personalized models”. In: Computers in Biology and Medicine 180 (2024), p. 108986. doi: 10.1016/j.compbiomed.2024.108986.

[30] F. L. Morales et al. “Noninvasive Electrical Mapping Compared with the Paced QRS Complex for Optimizing CRT Programmed Settings and Predicting Multidimensional Response”. In: Journal of Cardiovascular Translational Research (2023), pp. 1–13. doi: 10.1007/s12265-023-10418-1.

[31] D. J. Bivona et al. “Cardiac Magnetic Resonance, Electromechanical Activation, Kidney Function, and Natriuretic Peptides in Cardiac Resynchronization Therapy Upgrades”. In: Journal of Cardiovascular Development and Disease 10.10 (2023), p. 409. doi: 10.3390/jcdd10100409.

[32] M. D. Cerqueira, N. J. Weissman, V. Dilsizian, A. K. Jacobs, S. Kaul, W. K. Laskey, D. J. Pennell, J. A. Rumberger, T. Ryan, and M. S. Verani. “Standardized myocardial segmentation and nomenclature for tomographic imaging of the heart: a statement for healthcare professionals from the Cardiac Imaging Committee of the Council on Clinical Cardiology of the American Heart Association”. In: Circulation 105.4 (2002), pp. 539–542. doi: 10.1161/hc0402.102975.

[33] G. S. Stergiou, B. Alpert, S. Mieke, R. Asmar, N. Atkins, S. Eckert, G. Frick, B. Friedman, T. Graßl, T. Ichikawa, et al. “A Universal Standard for the Validation of Blood Pressure Measuring Devices: Association for the Advancement of Medical Instrumentation/European Society of Hypertension/International Organization for Standardization (AAMI/ESH/ISO) Collaboration Statement”. In: Hypertension 71.3 (2018), pp. 368–374. doi: 10.1161/HYPERTENSIONAHA.117.10237.

[34] R. Coulden and E. Sonnex. “Inter-observer variation in LV analysis in a dedicated CMR unit: the impact of audit and consensus guideline on reproducibility”. In: Journal of Cardiovascular Magnetic Resonance 16.Suppl 1 (2014), P372. doi: 10.1186/1532-429X-16-S1-P372.

[35] S. E. Luijnenburg, D. Robbers-Visser, A. Moelker, H. W. Vliegen, B. J. M. Mulder, and W. A. Helbing. “Intra-observer and interobserver variability of biventricular function, volumes and mass in patients with congenital heart disease measured by CMR imaging”. In: The International Journal of Cardiovascular Imaging 26 (2010), pp. 57–64. doi: 10.1007/s10554-009-9501-y.

[36] W. Bai, M. Sinclair, G. Tarroni, O. Oktay, M. Rajchl, G. Vaillant, A. M. Lee, N. Aung, E. Lukaschuk, M. M. Sanghvi, et al. “Automated cardiovascular magnetic resonance image analysis with fully convolutional networks”. In: Journal of Cardiovascular Magnetic Resonance 20 (2018), p. 65. doi: 10.1186/s12968-018-0471-x.

[37] F. Plappert, P. J. Oomen, C. E. Jones, E. Charitakis, L. O. Karlsson, P. G. Platonov, M. Wallman, and F. Sandberg. “Computational model of haemodynamics during atrial fibrillation”. In: The Journal of Physiology (2025). doi: 10.1113/jp289469.

[38] J. Lumens, T. Delhaas, B. Kirn, and T. Arts. “Three-Wall Segment (TriSeg) Model Describing Mechanics and Hemodynamics of Ventricular Interaction”. In: Annals of Biomedical Engineering 37.11 (2009), pp. 2234–2255. doi: 10.1007/s10439-009-9774-2.

[39] T. Arts, P. Bovendeerd, T. Delhaas, and F. Prinzen. “Modeling the relation between cardiac pump function and myofiber mechanics”. In: Journal of Biomechanics 36.5 (2003), pp. 731–736. doi: 10.1016/s0021-9290(02)00451-7.

[40] J. Walmsley, T. Arts, N. Derval, P. Bordachar, H. Cochet, S. Ploux, F. W. Prinzen, T. Delhaas, and J. Lumens. “Fast Simulation of Mechanical Heterogeneity in the Electrically Asynchronous Heart Using the MultiPatch Module”. In: PLOS Computational Biology 11.7 (2015), e1004284. doi: 10.1371/journal.pcbi.1004284.

[41] M. D. Morris. “Factorial Sampling Plans for Preliminary Computational Experiments”. In: Technometrics 33.2 (1991), pp. 161–174. doi: 10.1080/00401706.1991.10484804.

[42] U. Noè, A. Lazarus, H. Gao, V. Davies, B. Macdonald, K. Mangion, C. Berry, X. Luo, and D. Husmeier. “Gaussian process emulation to accelerate parameter estimation in a mechanical model of the left ventricle: a critical step towards clinical end-user relevance”. In: Journal of The Royal Society Interface 16.156 (2019), p. 20190114. doi: 10.1098/rsif.2019.0114.

[43] C. Rodero, S. Longobardi, C. Augustin, M. Strocchi, G. Plank, P. Lamata, and S. A. Niederer. “Calibration of Cohorts of Virtual Patient Heart Models Using Bayesian History Matching”. en. In: Annals of Biomedical Engineering 51.1 (Jan. 2023), pp. 241–252. doi: 10.1007/s10439-022-03095-9.

[44] M. Strocchi, C. M. Augustin, M. A. F. Gsell, C. A. Rinaldi, E. J. Vigmond, G. Plank, C. J. Oates, R. D. Wilkinson, and S. A. Niederer. “Integrating Imaging and Invasive Pressure Data into a Multiscale Whole-Heart Model”. In: Journal of Biomechanical Engineering 148.5 (2025). doi: 10.1115/1.4069497.

[45] S. Qian et al. “Developing cardiac digital twin populations powered by machine learning provides electrophysiological insights in conduction and repolarization”. In: Nature Cardiovascular Research 4.5 (2025), pp. 624–636. doi: 10.1038/s44161-025-00650-0.

[46] S. Longobardi, A. Lewalle, S. Coveney, I. Sjaastad, E. K. S. Espe, W. E. Louch, C. J. Musante, A. Sher, and S. A. Niederer. “Predicting left ventricular contractile function via Gaussian process emulation in aortic-banded rats”. en. In: Philosophical Transactions of the Royal Society A: Mathematical, Physical and Engineering Sciences 378.2173 (June 2020), p. 20190334. doi: 10.1098/rsta.2019.0334.

[47] M. Strocchi et al. “Cell to whole organ global sensitivity analysis on a four-chamber heart electromechanics model using Gaussian processes emulators”. en. In: PLOS Computational Biology 19.6 (June 2023). Ed. by K. Tsaneva-Atanasova, e1011257. doi: 10.1371/journal.pcbi.1011257.

[48] M. A. Stoffel, B. M. Li, K. Westerling, S. Arana, M. Balmus, E. Daub, and S. Niederer. “AutoEmulate: A Python package for semi-automated emulation”. In: Journal of Open Source Software 10.107 (2025), p. 7626. doi: 10.21105/joss.07626.

[49] C. E. Rasmussen and C. K. I. Williams. Gaussian Processes for Machine Learning. The MIT Press, 2005. doi: 10.7551/mitpress/3206.001.0001.

[50] H. Dou, S. Virtanen, N. Ravikumar, and A. F. Frangi. “A Generative Shape Compositional Framework to Synthesize Populations of Virtual Chimeras”. In: IEEE Transactions on Neural Networks and Learning Systems 36.3 (2025), pp. 4750–4764. doi: 10.1109/TNNLS.2024.3374121.

[51] N. V. Chawla, K. W. Bowyer, L. O. Hall, and W. P. Kegelmeyer. “SMOTE: Synthetic Minority Over-sampling Technique”. In: Journal of Artificial Intelligence Research 16 (2002), pp. 321–357. doi: 10.1613/jair.953.

[52] M. E. Johnson, L. M. Moore, and D. Ylvisaker. “Minimax and maximin distance designs”. In: Journal of Statistical Planning and Inference 26.2 (1990), pp. 131–148. doi: 10.1016/0378-3758(90)90122-B.

[53] T. F. Gonzalez. “Clustering to minimize the maximum intercluster distance”. In: Theoretical Computer Science 38 (1985), pp. 293–306. doi: 10.1016/0304-3975(85)90224-5.

[54] C. J. F. ter Braak and J. A. Vrugt. “Differential Evolution Markov Chain with snooker updater and fewer chains”. In: Statistics and Computing 18.4 (2008), pp. 435–446. doi: 10.1007/s11222-008-9104-9.

[55] M. D. Hoffman and A. Gelman. “The No-U-Turn Sampler: Adaptively Setting Path Lengths in Hamiltonian Monte Carlo”. In: Journal of Machine Learning Research 15 (2014), pp. 1593–1623. doi: 10.5555/2627435.2638586.

[56] A. Gelman and D. B. Rubin. “Inference from iterative simulation using multiple sequences”. In: Statistical Science 7.4 (1992), pp. 457–472.

[57] A. Vehtari, A. Gelman, D. Simpson, B. Carpenter, and P.-C. Bürkner. “Rank-normalization, folding, and localization: An improved *R̂* for assessing convergence of MCMC”. In: Bayesian Analysis 16.2 (2021), pp. 667–718.

[58] K. Menon, A. Zanoni, M. Khan, G. Geraci, K. Nieman, D. Schiavazzi, and A. Marsden. “Personalized and uncertainty-aware coronary hemodynamics simulations: From Bayesian estimation to improved multi-fidelity uncertainty quantification”. In: Computer Methods and Programs in Biomedicine 271 (2025), p. 108951. doi: 10.1016/j.cmpb.2025.108951.

[59] M. Mamajiwala, C. Corrado, C. Lanyon, S. Niederer, R. Wilkinson, and R. Clayton. “Rapid calibration of atrial electrophysiology models using Gaussian process emulators in the ensemble Kalman filter”. In: Scientific Reports (2025). doi: 10.1038/s41598-026-39948-9.

[60] M. Beetz et al. “Interpretable cardiac anatomy modeling using variational mesh autoencoders”. In: Frontiers in Cardiovascular Medicine 9 (2022), p. 983868. doi: 10.3389/fcvm.2022.983868.

[61] F. Gu et al. “Physiology-Informed Digital Twin-AI Framework Predicts Pacing Therapy Response in HFpEF”. In: medRxiv (2026), p. 2026.03.06.26347199. doi: 10.64898/2026.03.06.26347199.

[62] A. Ramos, F. Fenton, and E. Cherry. “Bayesian inference for fitting cardiac models to experiments: estimating parameter distributions using Hamiltonian Monte Carlo and approximate Bayesian computation”. In: Medical & Biological Engineering & Computing 61 (2022), pp. 75–95. doi: 10.1007/s11517-022-02685-y.

[63] A. Ramos, C. Herndon, F. Fenton, and E. Cherry. “Quantifying Distributions of Parameters for Cardiac Action Potential Models Using the Hamiltonian Monte Carlo Method”. In: 2021 Computing in Cardiology (CinC). Vol. 48. 2021, pp. 1–4. doi: 10.23919/cinc53138.2021.9662836.

[64] M. Salvador, F. Regazzoni, L. Dedè, and A. Quarteroni. “Fast and robust parameter estimation with uncertainty quantification for the cardiac function”. In: Computer Methods and Programs in Biomedicine 231 (2022), p. 107402. doi: 10.48550/arxiv.2210.03012.

[65] M. Spitieris and I. Steinsland. “Bayesian Calibration of imperfect computer models using Physics-informed priors”. In: ArXiv abs/2201.06463 (2022).

[66] M. Strocchi et al. “His-bundle and left bundle pacing with optimized atrioventricular delay achieve superior electrical synchrony over endocardial and epicardial pacing in left bundle branch block patients”. In: Heart Rhythm 17.11 (2020), pp. 1922–1929. doi: 10.1016/j.hrthm.2020.06.028.

[67] S. Khamzin, A. Dokuchaev, A. Bazhutina, T. Chumarnaya, S. Zubarev, T. Lyubimtseva, V. Lebedeva, D. Lebedev, V. Gurev, and O. Solovyova. “Machine Learning Prediction of Cardiac Resynchronisation Therapy Response From Combination of Clinical and Model-Driven Data”. en. In: Frontiers in Physiology 12 (Dec. 2021), p. 753282. doi: 10.3389/fphys.2021.753282.

[68] A. Haque, D. Stubbs, N. Hubig, F. Spinale, and W. Richardson. “Interpretable machine learning predicts cardiac resynchronization therapy responses from personalized biochemical and biomechanical features”. In: BMC Medical Informatics and Decision Making 22 (2022). doi: 10.1186/s12911-022-02015-0.

[69] W. Nazar, S. Szymanowicz, K. Nazar, D. Kaufmann, E. Wabich, R. Braun-Dullaeus, and L. Daniłowicz-Szymanowicz. “Artificial intelligence models in prediction of response to cardiac resynchronization therapy: a systematic review”. In: Heart Failure Reviews 29 (2023), pp. 133–150. doi: 10.1007/s10741-023-10357-8.

[70] P. Wouters et al. “Electrocardiogram-based deep learning improves outcome prediction following cardiac resynchronization therapy”. In: European Heart Journal 44 (2022), pp. 680–692. doi: 10.1093/eurheartj/ehac617.

[71] J. D. Dworkin, K. A. Linn, E. G. Teich, P. Zurn, R. T. Shinohara, and D. S. Bassett. “The extent and drivers of gender imbalance in neuroscience reference lists”. In: Nature Neuroscience 23.8 (2020), pp. 918–926. doi: 10.1038/s41593-020-0658-y.

[72] B. Rowson, S. M. Duma, M. R. King, I. Efimov, A. Saterbak, and N. C. Chesler. “Citation Diversity Statement in BMES Journals”. In: Annals of Biomedical Engineering 49.3 (2021), pp. 947–949. doi: 10.1007/s10439-021-02739-6.

[73] D. Zhou, E. J. Cornblath, J. Stiso, E. G. Teich, J. D. Dworkin, A. S. Blevins, and D. S. Bassett. Gender Diversity Statement and Code Notebook v1.0. 2020. doi: 10.5281/zenodo.3672110.

[74] I. Van Buskirk, A. Clauset, and D. B. Larremore. “An Open-Source Cultural Consensus Approach to Name-Based Gender Classification”. In: Proceedings of the International AAAI Conference on Web and Social Media. Vol. 17. 2023, pp. 866–877. doi: 10.1609/icwsm.v17i1.22195.

[75] S. Laohaprapanon, G. Sood, and B. Naji. ethnicolr: Predict Race and Ethnicity From Name. https://github.com/appeler/ethnicolr. Version 0.9.8. 2022.

[76] J. N. Cohn, T. B. Levine, M. T. Olivari, V. Garberg, D. Lura, G. S. Francis, A. B. Simon, and T. Rector. “Plasma Norepinephrine as a Guide to Prognosis in Patients with Chronic Congestive Heart Failure”. In: New England Journal of Medicine 311.13 (1984), pp. 819–823. doi: 10.1056/NEJM198409273111303.

[77] R. J. Cody, G. J. Haas, P. F. Binkley, Q. Capers, and R. Kelley. “Plasma endothelin correlates with the extent of pulmonary hypertension in patients with chronic congestive heart failure”. In: Circulation 85.2 (1992), pp. 504–509. doi: 10.1161/01.CIR.85.2.504.

[78] S. S. Gottlieb, K. Dickstein, E. Fleck, J. Kostis, T. B. Levine, T. LeJemtel, and M. DeKock. “Hemodynamic and neurohormonal effects of the angiotensin II antagonist losartan in patients with congestive heart failure”. In: Circulation 88.4 (1993), pp. 1602–1609. doi: 10.1161/01.CIR.88.4.1602.

[79] R. Pacher, J. Bergler-Klein, S. Globits, H. Teufelsbauer, M. Schuller, A. Krauter, E. Ogris, S. Rödler, M. Wutte, and E. Hartter. “Plasma big endothelin-1 concentrations in congestive heart failure patients with or without systemic hypertension”. In: The American Journal of Cardiology 71.15 (1993), pp. 1293–1299. doi: 10.1016/0002-9149(93)90543-L.

[80] M. Nootens, E. Kaufmann, T. Rector, C. Toher, D. Judd, G. S. Francis, and S. Rich. “Neurohormonal activation in patients with right ventricular failure from pulmonary hypertension: Relation to hemodynamic variables and endothelin levels”. In: Journal of the American College of Cardiology 26.7 (1995), pp. 1581–1585. doi: 10.1016/0735-1097(95)00399-1.

[81] T. Tsutamoto, T. Hisanaga, A. Wada, K. Maeda, M. Ohnishi, D. Fukai, N. Mabuchi, M. Sawaki, and M. Kinoshita. “Interleukin-6 Spillover in the Peripheral Circulation Increases With the Severity of Heart Failure, and the High Plasma Level of Interleukin-6 Is an Important Prognostic Predictor in Patients With Congestive Heart Failure”. In: Journal of the American College of Cardiology 31.2 (1998), pp. 391–398. doi: 10.1016/S0735-1097(97)00494-4.

