## Supplemental Material for "Population-level emulation enables fast Bayesian calibration of cardiac digital twins"

*Annals of Biomedical Engineering*

### S1 CONSTRUCTION OF DIGITAL CHIMERAS

This section expands on the digital chimeras of the main text (Section 2.3.3). Patients are represented by their latent encodings $\mathcal{L}^{(i)} \in \mathbb{R}^k$, obtained by standardizing the patient features $\mathcal{F}^{(i)}$ and projecting them onto the $k$ leading principal components (main text, Section 2.3.1).

**Convex combination.** A chimera is formed from two members $i, j$ of the current pool, which holds the real patients and all previously accepted chimeras, as

$$\tilde{\mathcal{L}} = \alpha \mathcal{L}_i + (1 - \alpha)\mathcal{L}_j. \quad \text{(S1)}$$

We used $\alpha = 0.5$, so that both parents contribute equally, although any $\alpha \in [0, 1]$ may be chosen. Because the projection from features to latent space is affine, applying the same convex combination to the continuous patient features reproduces Eq. (S1) exactly in latent space.

**Simulator inputs.** We use this property to construct each chimera's simulator inputs. Continuous scalar and array-valued simulator inputs (heart rate, body surface area, atrioventricular delay, segmental activation times and wall volumes) are interpolated element-wise. Ischemia indicators are kept binary rather than averaged: every binary combination of the segments on which the parents disagree is generated as a separate candidate, and the greedy selection below chooses among them. Non-numeric fields, such as the simulation input template, are inherited from the parent nearest the chimera in latent space. Because the binary candidates lie off the interpolated point, the emulator receives the latent encoding of each chimera's actual simulator inputs, obtained by projecting those inputs with the same standardization and principal components, rather than Eq. (S1). Apart from the ischemia indicators, every chimera is a convex combination of real patients, so the augmentation densifies the interior of the sampled region without extending it.

**Recursive greedy-maximin selection.** We select $\tilde{N}$ chimeras with an incremental greedy-maximin procedure. The candidates are initially the $\binom{N}{2}$ midpoints between pairs of real patients. At each iteration, the candidate with the greatest minimum distance to every current anchor (all real patients and all previously selected chimeras) is accepted and added to the anchor set. The candidate pool then grows rather than only shrinking: every midpoint between the newly accepted chimera and each existing anchor is added as a new candidate, so the space recursively subdivides. If two nearby chimeras are selected in successive rounds, a later round can propose the point between them and close the gap. With a static pool that only removes candidates, coverage would plateau once the initial $\binom{N}{2}$ midpoints are exhausted; with the growing pool it keeps improving with the chimera count.

**Pruning.** Because the pool can grow to tens of thousands of live candidates when hundreds of chimeras are selected, we prune it. For a candidate formed from two anchors $\mathcal{L}_i$ and $\mathcal{L}_j$, the minimum distance to the anchor set is at most $\min(\alpha, 1 - \alpha)\,\|\mathcal{L}_i - \mathcal{L}_j\|$, because both parents are anchors, and adding anchors can only reduce it further. Once this bound falls at or below the best minimum distance among the live candidates, the candidate cannot win the current selection and is discarded. Candidates with re-assigned ischemia indicators have no such bound and are never pruned. In practice this eliminates the large majority of candidates after each selection, keeping the growing pool tractable for chimera counts in the hundreds to low thousands.

### S2 PHYSIOLOGICAL BOUNDS FOR TRAINING SIMULATIONS

A training simulation was discarded if MONARCH did not reach a periodic steady state (limit cycle), defined by a relative difference in chamber volumes between consecutive beats below 0.01, or if any of its ten outputs fell outside the bounds in Table S1. For the 128-sample pool used by the calibration emulators, this discarded 34.4% of the simulations (main text, Section 2.3.2).

**Table S1:** Physiological bounds on the simulated outputs used to discard training simulations.

| Output | Description | Lower | Upper | Unit |
|---|---|---|---|---|
| LVEDV | LV end-diastolic volume | 30 | 600 | mL |
| LVESV | LV end-systolic volume | 5 | 550 | mL |
| SBP | Systolic blood pressure | 50 | 260 | mmHg |
| DBP | Diastolic blood pressure | 20 | 160 | mmHg |
| RVEDV | RV end-diastolic volume | 15 | 550 | mL |
| EDWthLfw | End-diastolic LFW thickness | 2 | 22 | mm |
| EDWthSw | End-diastolic SW thickness | 2 | 22 | mm |
| LVPER | LV peak ejection rate | 30 | 3,000 | $mL\,s^{-1}$ |
| LVCO | LV cardiac output | 0.3 | 13 | $L\,min^{-1}$ |
| LVEDP | LV end-diastolic pressure | 2 | 60 | mmHg |

MONARCH limit-cycle convergence: relative volume difference between consecutive beats $< 0.01$.

### S3 MCMC CONVERGENCE DIAGNOSTICS

Table S2 lists the convergence diagnostics of every calibration: the held-out calibrations with differential evolution Metropolis variant (DEMetropolisZ) (4 chains of 20,000 draws after 10,000 tuning steps) , reported in the main text. $\hat{R}$ is the rank-normalized split-$\hat{R}$ and ESS the effective sample size, computed over all chains with ArviZ; for each patient the table gives the worst value over the seven parameters. 21 of the 26 held-out calibrations meet both thresholds; the other five (patients 6, 10, 14, 15 and 20) exceed $\hat{R} = 1.01$ by at most 0.010, and patient 20 also has a bulk effective sample size just below 400.

Figs. S1–S3 show the held-out DEMetropolisZ chains of three patients: patient 10, the least well converged (highest $\hat{R}$); patient 8, typical of the cohort (its maximum $\hat{R}$ equals the cohort median); and patient 3, the patient most isolated in latent space, for whom only 33 of 100 posterior draws produced a valid MONARCH simulation (main text, Section 3.3). The tuning draws were not stored, so the traces begin after burn-in. For legibility, every tenth draw is plotted (2,000 per chain); the diagnostics use all draws. Trace axes and density axes span the uniform prior of each parameter, so the width of each density relative to the axis shows how strongly the clinical targets constrain that parameter.

**Table S2:** Convergence diagnostics for all 26 patients. Held-out calibrations with DEMetropolisZ. For each patient, the maximum $\hat{R}$ and the minimum bulk and tail effective sample sizes (ESS) over the seven parameters; bold marks values that miss $\hat{R} \leq 1.01$ or ESS $\geq 400$. $t$, wall-clock time of the held-out calibration on one CPU core; Valid, posterior draws (of 100) that produced a valid MONARCH simulation in the held-out posterior predictive check; Isch., at least one ischemic segment.

| | | Held-out DEMetropolisZ | | | | |
|---|---|---|---|---|---|---|
| Pt. | Isch. | $\hat{R}_{\max}$ | $\mathrm{ESS}_{\min}^{\mathrm{bulk}}$ | $\mathrm{ESS}_{\min}^{\mathrm{tail}}$ | $t$ (s) | Valid (/100) |
| 1 | | 1.004 | 1,761 | 1,275 | 61 | 97 |
| 2 | ✓ | 1.006 | 1,321 | 1,334 | 60 | 97 |
| 3 | ✓ | 1.003 | 2,536 | 2,118 | 62 | 33 |
| 4 | | 1.003 | 2,228 | 2,150 | 61 | 79 |
| 5 | | 1.007 | 776 | 685 | 61 | 100 |
| 6 | | **1.015** | 453 | **338** | 63 | 99 |
| 7 | | 1.003 | 1,860 | 1,531 | 62 | 98 |
| 8 | | 1.005 | 1,467 | 1,154 | 60 | 99 |
| 9 | | 1.003 | 1,665 | 995 | 62 | 98 |
| 10 | | **1.020** | 533 | **317** | 60 | 95 |
| 11 | | 1.004 | 1,190 | 1,035 | 62 | 59 |
| 12 | | 1.005 | 757 | 974 | 61 | 46 |
| 13 | | 1.003 | 1,156 | 1,304 | 60 | 100 |
| 14 | | **1.015** | 477 | **192** | 61 | 96 |
| 15 | | **1.014** | 487 | **163** | 61 | 25 |
| 16 | | 1.003 | 1,120 | 617 | 60 | 99 |
| 17 | ✓ | 1.006 | 852 | 487 | 61 | 98 |
| 18 | | 1.003 | 1,633 | 1,580 | 60 | 78 |
| 19 | | 1.007 | 1,339 | 1,443 | 62 | 100 |
| 20 | | **1.018** | **372** | 405 | 60 | 85 |
| 21 | ✓ | 1.002 | 2,242 | 2,497 | 61 | 97 |
| 22 | | 1.006 | 1,802 | 1,487 | 62 | 99 |
| 23 | ✓ | 1.002 | 2,147 | 2,216 | 62 | 92 |
| 24 | ✓ | 1.004 | 1,258 | 1,114 | 61 | 100 |
| 25 | | 1.006 | 1,733 | 1,291 | 61 | 97 |
| 26 | | 1.005 | 1,493 | 1,003 | 61 | 94 |

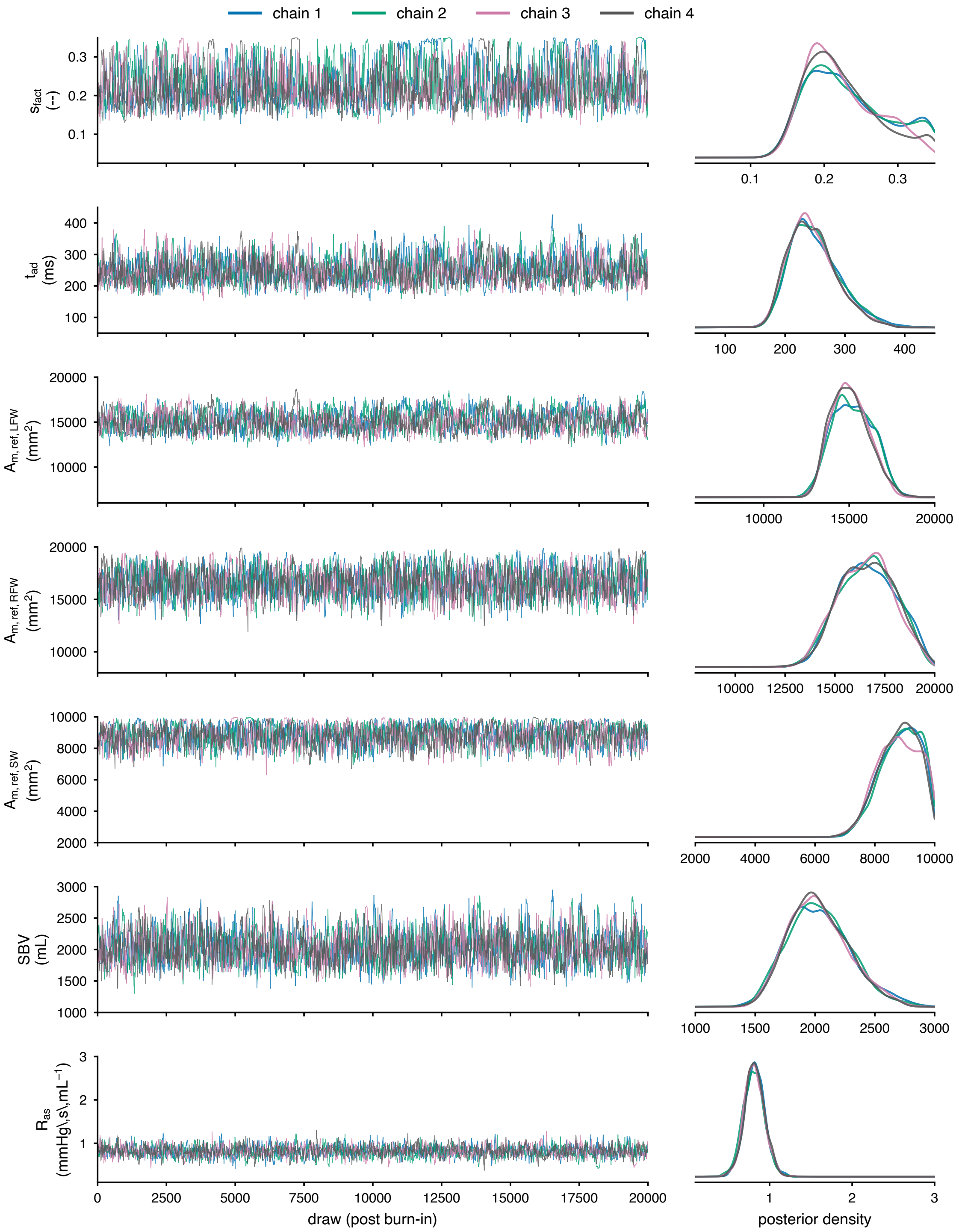


**Figure S1: Held-out DEMetropolisZ chains for patient 10, the least well converged calibration** (maximum $\hat{R}$ 1.020, minimum bulk ESS 533). Left: traces of the four chains after burn-in (every tenth draw); right: the posterior density of each chain. Rows show the seven free parameters (main text, Table 2).

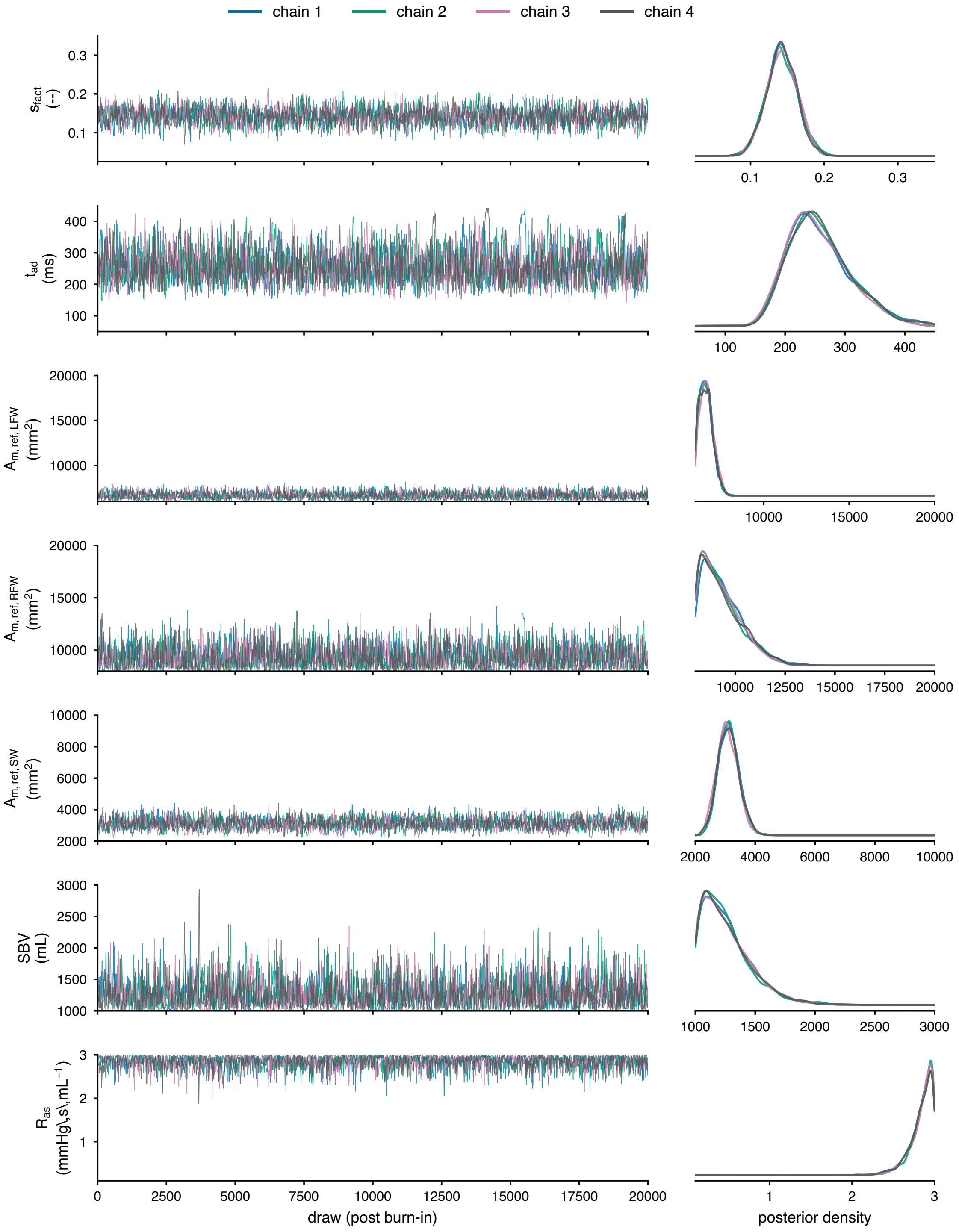


**Figure S2: Held-out DEMetropolisZ chains for patient 8, a typical calibration** (maximum $\hat{R}$ 1.005, minimum bulk ESS 1,467). Layout as in Fig. S1.

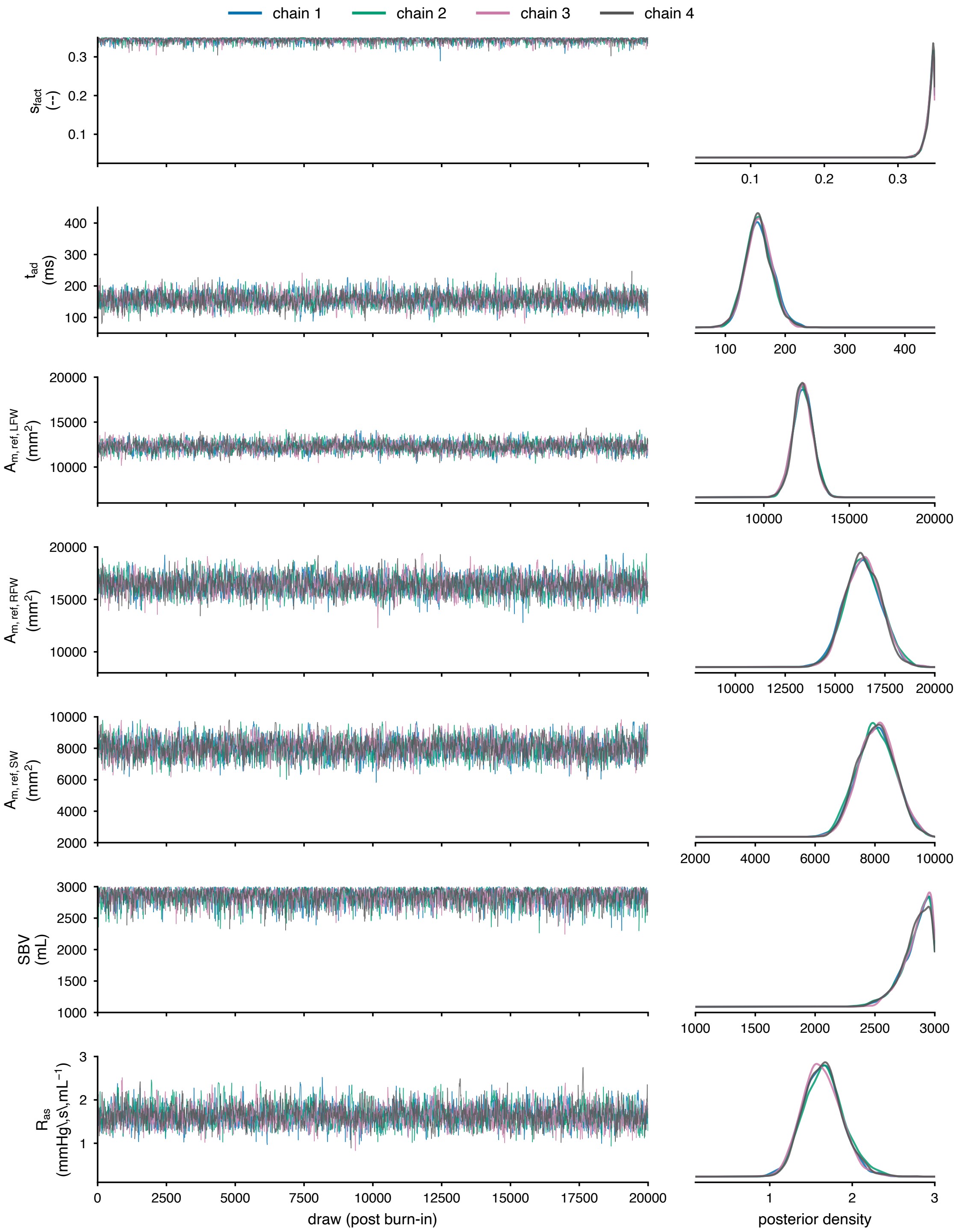


**Figure S3: Held-out DEMetropolisZ chains for patient 3, the patient most isolated in latent space** (maximum $\hat{R}$ 1.003, minimum bulk ESS 2,536). The chains converge although only 33 of 100 posterior draws produced a valid MONARCH simulation. Layout as in Fig. S1.

### S4 PARAMETER IDENTIFIABILITY

Fig. S4 shows the posterior contraction, $1 - \sigma_{\text{post}}/\sigma_{\text{prior}}$, of each free parameter for each patient, from the held-out calibrations (main text, Section 2.5). Mean contraction ranged from 0.57 for SBV to 0.83 for $A_{m,\text{ref,LFW}}$, and SBV was poorly constrained in a few patients. Because contraction is measured relative to the width of the uniform prior, it depends on the prior range: a parameter with a narrow prior can appear weakly constrained even when the data determine it well. The correlations between parameters in the posterior complement it (Fig. S5). The strongest, between $A_{m,\text{ref,LFW}}$ and $A_{m,\text{ref,SW}}$ (mean $|r| = 0.60$), was negative on average (mean $r = -0.54$): a larger left free wall was offset by a smaller septum, consistent with both walls jointly setting the left-ventricular cavity size. All other pairs had mean $|r| \leq 0.44$.

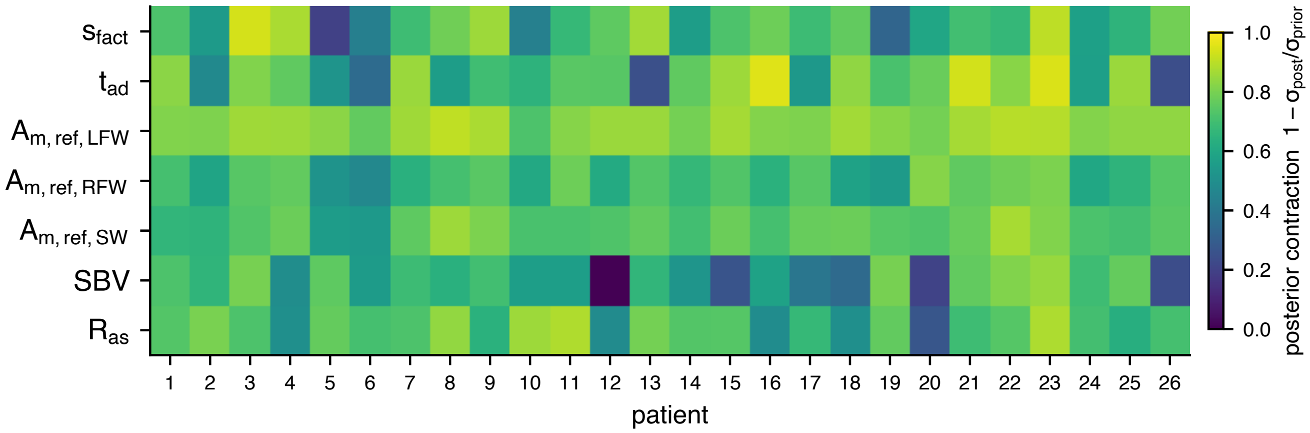


**Figure S4: Posterior contraction of the seven free parameters.** Posterior contraction, $1 - \sigma_{\text{post}}/\sigma_{\text{prior}}$, of each free parameter (rows) for each patient (columns), from the held-out calibrations; 0 means the posterior is as wide as the uniform prior and 1 that it collapses to a point. Parameters as in Table 2 of the main text.

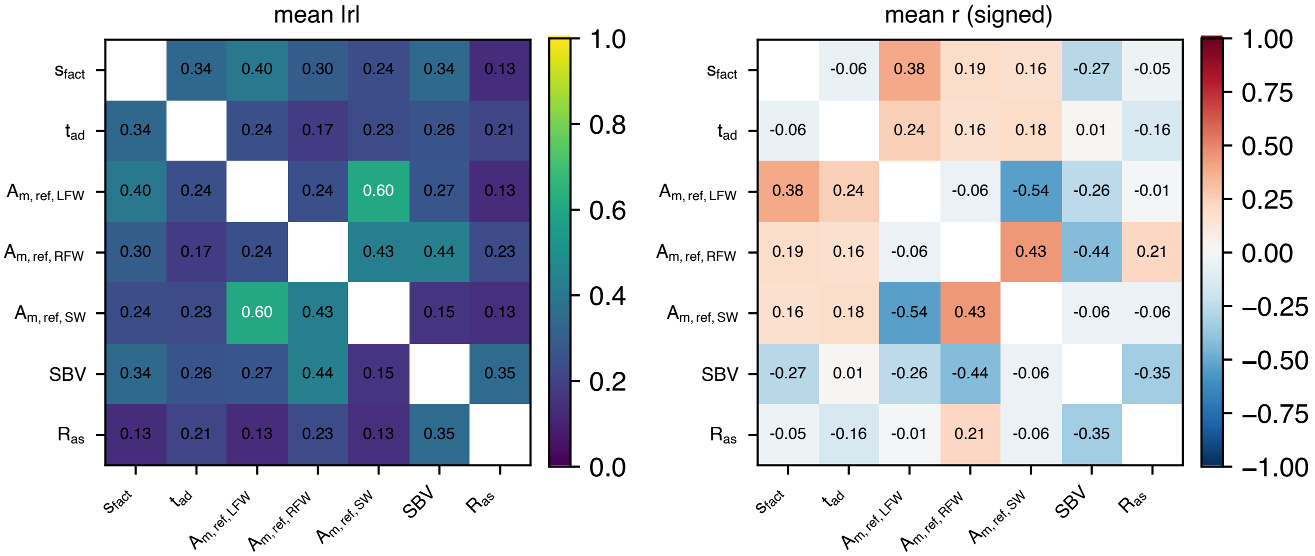


**Figure S5: Posterior correlations between the seven free parameters.** Pearson correlation between each pair of parameters in each patient's held-out posterior, averaged over the 26 patients. Left: mean absolute correlation $|r|$. Right: mean signed correlation $r$; negative values indicate a trade-off, in which the data constrain a combination of the two parameters more tightly than either alone. Parameters as in Table 2 of the main text.
